# Local glycolysis supports injury-induced axonal regeneration

**DOI:** 10.1101/2024.01.29.577706

**Authors:** Luca Masin, Steven Bergmans, Annelies Van Dyck, Lies De Groef, Karl Farrow, Lieve Moons

## Abstract

Successful axonal regeneration following injury requires the effective allocation of energy. Mitochondria play a pivotal role, accumulating at the tips of growing axons to fuel regeneration. However, how axons withstand the initial disruption in mitochondrial energy production caused by the injury, and subsequently initiate regrowth is poorly understood. Using retinal cultures in a multicompartment microfluidic device, we observed increased regrowth and enhanced mitochondrial trafficking in the axons of retinal ganglion cells with the deletion of both *Pten* and *Socs3*. While wild-type axons relied on mitochondrial metabolism, after injury, in the absence of *Pten* and *Socs3*, energy production was demonstrated to be supported by local glycolysis. Slowing down glycolysis in these axons impaired both regrowth and energy production. Together, these observations reveal that glycolytic ATP, combined with sustained mitochondrial transport, is essential for injury-induced axonal regrowth, providing new insights into the metabolic underpinnings of axonal regeneration.

## INTRODUCTION

The adult mammalian central nervous system (CNS) has a very limited regenerative capacity. Upon injury, neurons fail to regenerate their axons and often die, leaving irreversible damage^1,2^. Regrowth is limited by a wide array of both neuron-intrinsic mechanisms, including the progressive reduction in expression of regeneration-associated genes, as well as neuron-extrinsic ones, such as the presence of growth-inhibiting molecules and glial scarring^1,2^. Despite decades of research and the discovery of regeneration models, such as the genetic deletion of *Pten* and *Socs3*^3,4^, full circuit restoration and functional recovery remain elusive.

Axonal regeneration is an extremely energy demanding process and over the years mitochondria have emerged as critical players. The transport of functional mitochondria to the axons and their accumulation at the growth cone is crucial for regeneration^5–9^. However, injury has been reported to depolarize the local population of mitochondria in axons^7,10^ and sustained axonal ATP depletion is one of the main causes of Wallerian degeneration^11,12^. Unfortunately, mitochondria are predominantly stationary in the adult mammalian CNS, localizing at the site of synapses to provide the energy required for firing and neurotransmitter release^13^. As such, while remobilization of mitochondria is a viable strategy to sustain axonal regeneration, axons need to first withstand the injury, in the presence of a disrupted cytosolic environment and local dysfunctional mitochondria before the reallocation of healthy mitochondria from other neuronal compartments via biogenesis and trafficking is complete. Therefore, mitochondria-independent mechanisms might be at play after injury and might be required to initiate axonal regeneration.

Using an in vitro culture of postnatal retinal cells in microfluidic devices, this study shows that injury-induced axonal regeneration of retinal ganglion cells (RGCs) is associated with mitochondrial transport within the axons and that the increased axonal regrowth phenotype observed upon deletion of *Pten* and *Socs3* in RGCs is accompanied by enhanced mitochondrial trafficking. Nevertheless, we found that *Pten*^-/-^;*Socs3^-/-^*neurons upregulate glycolysis locally in the axonal compartment immediately after axotomy. Downregulation of glycolysis via galactose treatment completely abolishes the enhanced regeneration phenotype of *Pten*^-/-^;*Socs3^-/-^* axons and impairs their ability to produce ATP. This work demonstrates the importance axonal glycolysis plays as a source of energy during the initial phases of axonal regrowth.

## RESULTS

### Deletion of *Pten* and *Socs3* increases axonal regeneration of RGCs *in vitro*

To study the metabolism underlying axonal regeneration we established a culture of primary retinal cells from postnatal *Pten^fl/fl^*;*Socs3^fl/fl^*mice in microfluidic devices. Multicompartment microfluidics allowed us to separate the neuronal cell bodies in the somatic channel from their axons, which can grow through the microgrooves and extend into the axonal channels^14^. This fluidic isolation enables clean axonal injuries and independent treatment of the axonal and somato-dendritic compartments. After two weeks in culture, we identified retinal ganglion cells in the somatic channel using Tuj1 and RBPMS immunolabelling, both RGC markers in the murine retina^15,16^(Fig S1A). Dendrites were restricted to the somatic channel, while many axons extended into the axonal channel, as revealed by Map2 and Tuj1 labelling, respectively (Fig S1B). As no purification step was carried out during isolation, we could also identify retinal astrocytes and Müller glia in the somatic channel via GFAP and GLAST labelling (Fig S1C).

To elucidate the mechanisms underlying the regeneration induced by *Pten* and *Socs3* deletion, we divided the retinal cultures into two conditions: *Pten^fl/fl^*;*Socs3^fl/fl^* cultures with no induced recombination, here called wild-type (WT), and *Pten^fl/fl^*;*Socs3^fl/fl^* cultures, in which we induced the conditional double-knockout of *Pten* and *Socs3* (cdKO) specifically in RGCs via AAV-mediated Cre recombination (*AAV2/2-hSyn1-Cre-t2A-mKate2*) (Fig 1A). At 14 days in vitro, PTEN immunolabeling was not detectable in 98% of the RGC somata, which were co-labelled by hSyn-YFP and Tuj1 (Fig S1D-E), highlighting efficient and specific recombination.

**Figure 1:**
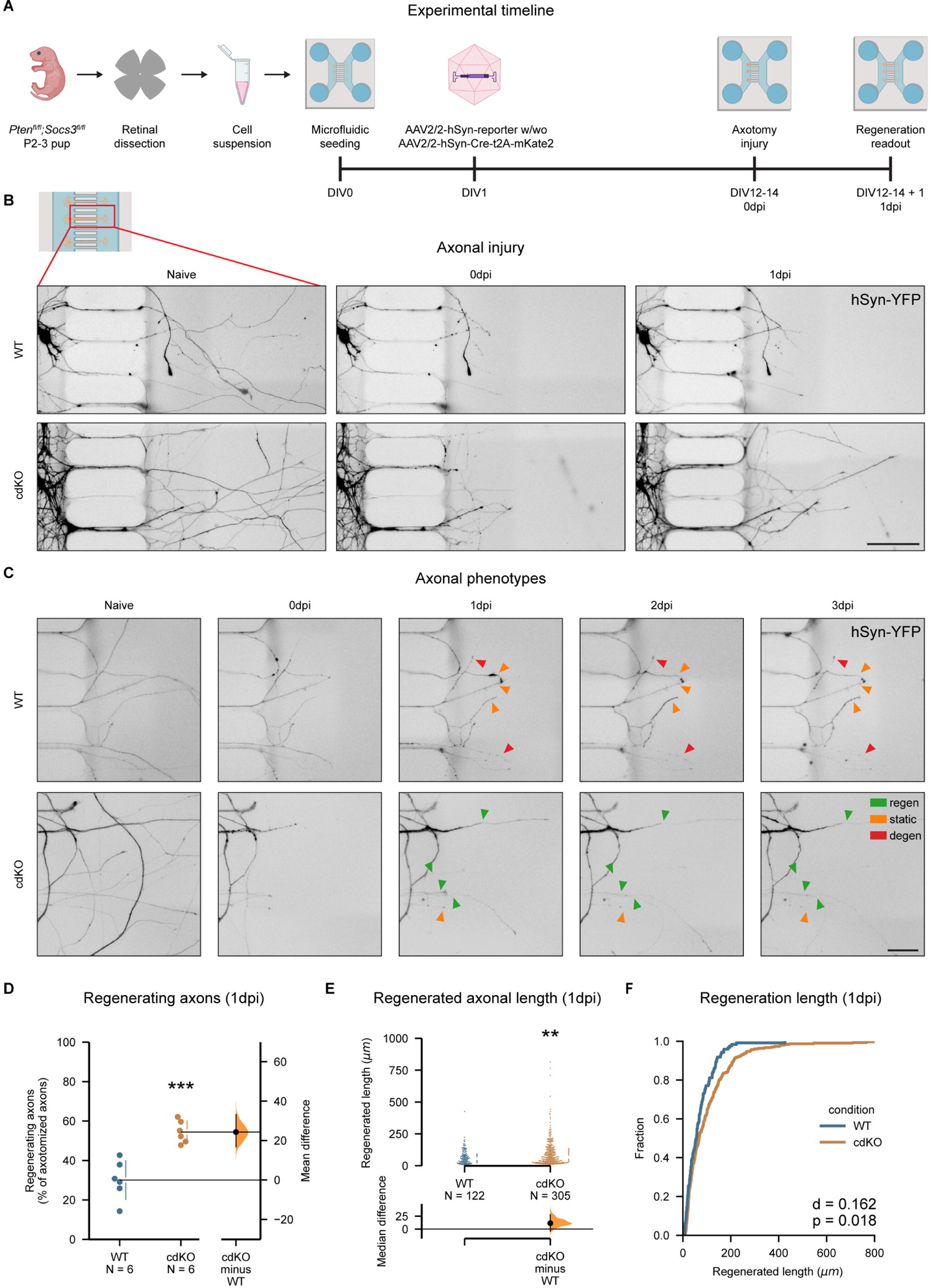
Deletion of Pten and Socs3 increases axotomy-induced axonal regeneration in primary RGCs. (A) Primary retinal cultures are obtained from retinae of *Pten^fl/fl^;Socs3^fl/fl^* mice at P2-3 and seeded in microfluidic devices. The RGCs are labelled and conditionally deleted for *Pten* and *Socs3* via AAV2/2-mediated transduction at 1 day *in vitro* (DIV1). Axonal injury is performed between DIV12 and DIV14 and the outcome of regeneration is assessed one day later (1dpi, DIV12-14+1). (B) Representative images of uninjured axons (naive), injured axons immediately after axotomy (0dpi) and regrowing axons (1dpi) of WT and cdKO RGCs. hSyn-YFP is encoded by an AAV2/2-hSyn-ATeam^YEMK^-WPRE-hGHp vector (see Methods). Scale bar 100 µm. (C) Representative images of degenerating axons (red), regenerating axons (green) and static axons (orange). The latter survive injury but do not grow past the cut site at 1dpi and until 3dpi. Scale bar 50 µm. (D) Quantification of the axons regrowing in the axonal compartment shows that deletion of *Pten* and *Socs3* induces an increase in the percentage of regenerating axons at 1dpi. (E) Quantification of the axonal length shows that deletion of *Pten* and *Socs3* induces a significant increase in the average length of regrowing axons at 1dpi. (F) Cumulative distribution of axonal lengths shows that *Pten* and *Socs3* deletion increases the length of regrowth at 1dpi. Data from 6 independent experiments, presented as mean ± SD (C) or median ± 25-75^th^CI (E) and bootstrap 95% CI versus WT. Student t-test (D), Kolmogorov–Smirnov test (E) and Mann-Whitney U-test (F). **p<0.01, ***p<0.001. See also Fig S1.

To quantify the regeneration capacity of WT and cdKO RGCs, we performed an axotomy in the axonal channel, and classified the axons in 3 groups one day after injury (1 dpi) (Fig 1C). The three classes were: 1) regenerating axons, which extended past the site of injury, 2) degenerating axons, which either fully degenerated or showed substantial beading and rupture and 3) static axons, which showed minimal or no sign of beading, but failed to extend past the site of injury at 1dpi. Of note, when followed up until 3dpi, axons classified as static on 1dpi failed to show either signs of degeneration or growth past the site of injury (Fig 1C). The relative percentages of these axon classes varied upon co-deletion of *Pten* and *Socs3* (Fig S1F), with a significant increase in the percentage of regenerating axons at 1dpi for cdKO RGCs as compared to WT (Fig 1D). We also detected an increase in the median length of the regrown axon tracts at 1dpi in cdKO RGCs as compared to WT neurons (Fig 1E-F). Finally, besides the increased regeneration, the deletion of *Pten* and *Socs3* led to a decrease in the percentage of degenerating axons at 1dpi as compared to WT (Fig S1G). Overall, these findings show that deletion of *Pten* and *Socs3* enhances axonal regeneration in cultured RGCs.

### Axonal regeneration is associated with mitochondrial transport and integrity

Mitochondrial transport into the axons has been shown to be important for regeneration in other models^7^ and deletion of *Pten* and *Socs3* was reported to be associated with increased axonal transport of mitochondria in embryonic cortical neurons^6^. As we hypothesized this to hold true for cultured postnatal RGCs, we transduced them with a vector encoding a mitochondrially-targeted GFP to label the mitochondria, as well as a cytosolic mCherry to visualize the whole neuron (*AAV2/2-CAG-FLeX-mitoGFP-t2A-mCherry-WPRE*) (Fig 2A). We then quantified the transport of mitochondria via kymographic analysis (Fig 2B) in uninjured axons (naive), within the first hour after injury (0dpi) and one day after injury (1dpi). Axonal injury caused a disruption of mitochondrial transport in both WT and cdKO axons, and trafficking was not restored in static axons at 1dpi (Fig 2C-D). Regenerating WT axons displayed transport comparable to their uninjured counterparts, while cdKO regenerating axons showed upregulated trafficking compared to naive (Fig 2D) and compared to regenerating WT axons (Fig S2A). A similar genotypic difference in regenerating axons was detected when measuring anterograde transport only (Fig S2B).

**Figure 2:**
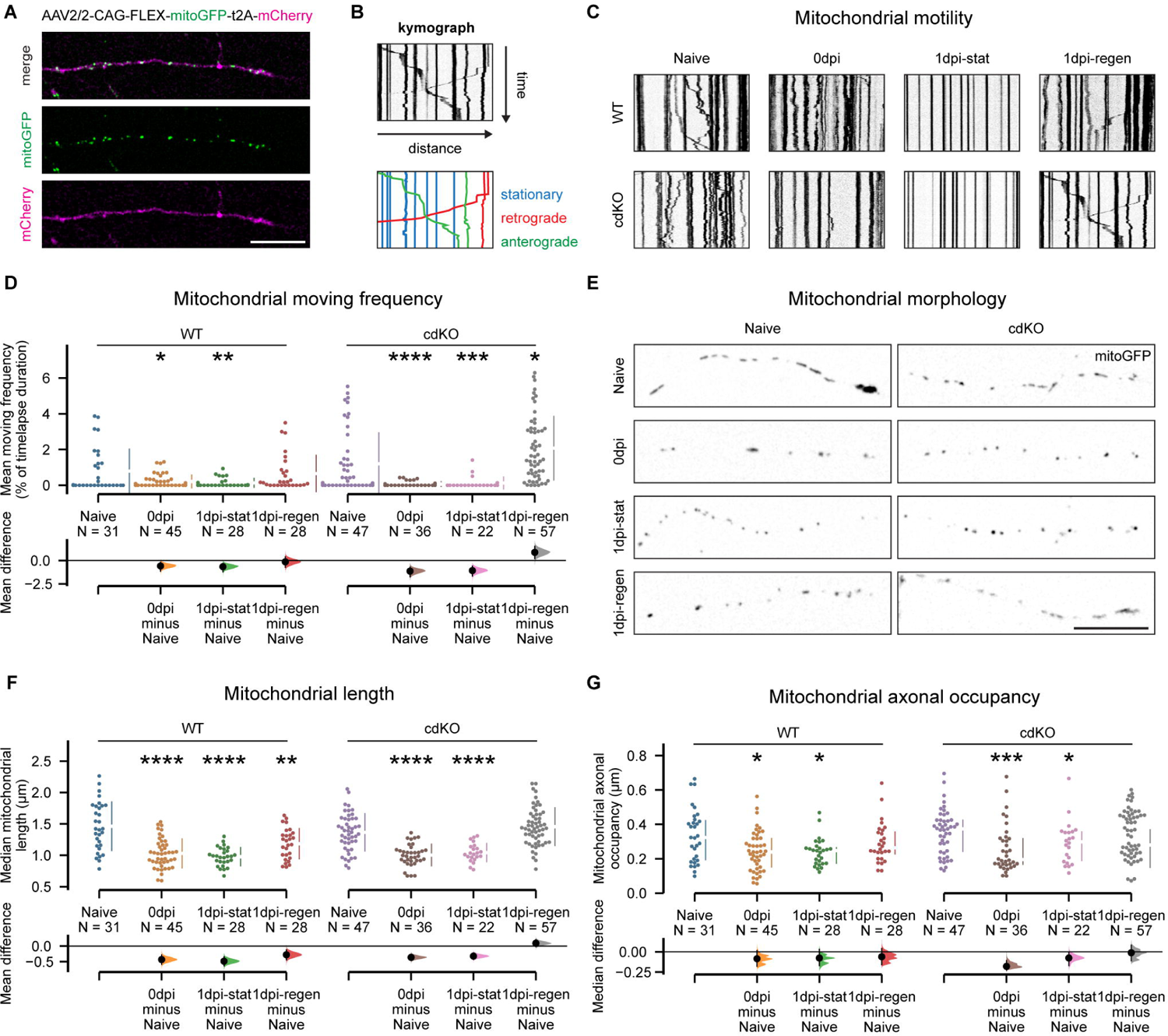
Axonal regeneration is associated with the restoration of mitochondrial transport, morphology, and mass. (A) RGC axons and their mitochondria are labelled via viral expression of mCherry and a mitochondrially-targeted GFP, respectively. Scale bar 25 um. (B) Example kymograph highlighting mitochondria which are classified as stationary (blue), transported retrogradely (red) and anterogradely (green). (C) Representative kymographs showing mitochondrial transport for WT and cdKO axons, both in naive and injured conditions. (D) Quantification of mitochondrial moving frequency per axon reveals that regenerating axons show a similar (WT) or higher (cdKO) mitochondrial transport as compared to naive. (E) Representative images of mitochondrial morphology in WT and cdKO axons, showing small round mitochondria immediately after injury and in static axons. Scale bar 20 µm (F) Quantification of the average mitochondrial length per axon shows full restoration to lengths comparable with naive mitochondria only in regenerating cdKO axons. (G) Quantification of mitochondrial axonal occupancy shows recovery of mitochondrial mass in regenerating axons, but not in static ones, for both WT and cdKO RGCs. Data from 4 independent experiments, presented as mean ± SD (D, F) or median ± 25-75^th^CI (G) and bootstrap 95% CI versus relative naive. Welch (D, F) and Kruskal-Wallis ANOVA (G). *p<0.05, **p<0.01, ***p<0.001, ****p<0.0001. See also Fig S2.

Furthermore, axonal injury caused a significant decrease in mitochondrial length in both WT and cdKO axons at 0dpi, leading to small, round mitochondria, possibly due to injury-induced fission (Fig 2E-F). At 1dpi, only cdKO regenerating axons were able to fully recover mitochondrial length (Fig 2F), which was significantly higher than in regenerating WT axons (Fig S2C). On the other hand, static axons of both WT and cdKO RGCs did not restore an elongated mitochondrial morphology comparable to naive (Fig 2E-F). Lastly, as a measure of mitochondrial mass, we calculated the mitochondrial axonal occupancy^17^. As for mitochondrial length, we measured a small but significant decrease in occupancy after injury at 0dpi, which was not restored at 1dpi in static axons of both WT and cdKO RGCs. Regenerating axons on the other hand, restored occupancy to a value comparable to their naive counterparts in both genotypes. (Fig 2F). Of note, no significant difference in mitochondrial transport, morphology and occupancy was measured in uninjured axons between WT and cdKO RGCs (Fig S2A-D). Taken together, these data shows that axonal regeneration is associated with mitochondrial transport and integrity and the latter are enhanced after injury by deletion of *Pten* and *Socs3*.

### Preservation of energy levels in cdKO axons is supported by glycolysis

To determine whether the differences in mitochondrial phenotype between static and regenerating axons were reflected in the axonal energy levels, we expressed the ATeam1.03^YEMK^ biosensor^18^ (*AAV2/2-hSyn1-ATeam^YEMK^-WPRE-hGHp*) and measured the intra-axonal concentration of ATP. Consistent with previous studies^19^, injury caused a significant decrease in axonal ATP at 0dpi, in both WT and cdKO axons (Fig 3A-B). At 1dpi, in WT RGCs, only regenerating axons restored ATP to levels comparable with naive. On the other hand, we detected an ATP concentration comparable to naive in cdKO static axons and a significantly higher ATP concentration as compared to naive in cdKO regenerating axons (Fig 3A-B). No significant difference in ATP was measured between WT and cdKO uninjured axons (Fig S3A).

**Figure 3:**
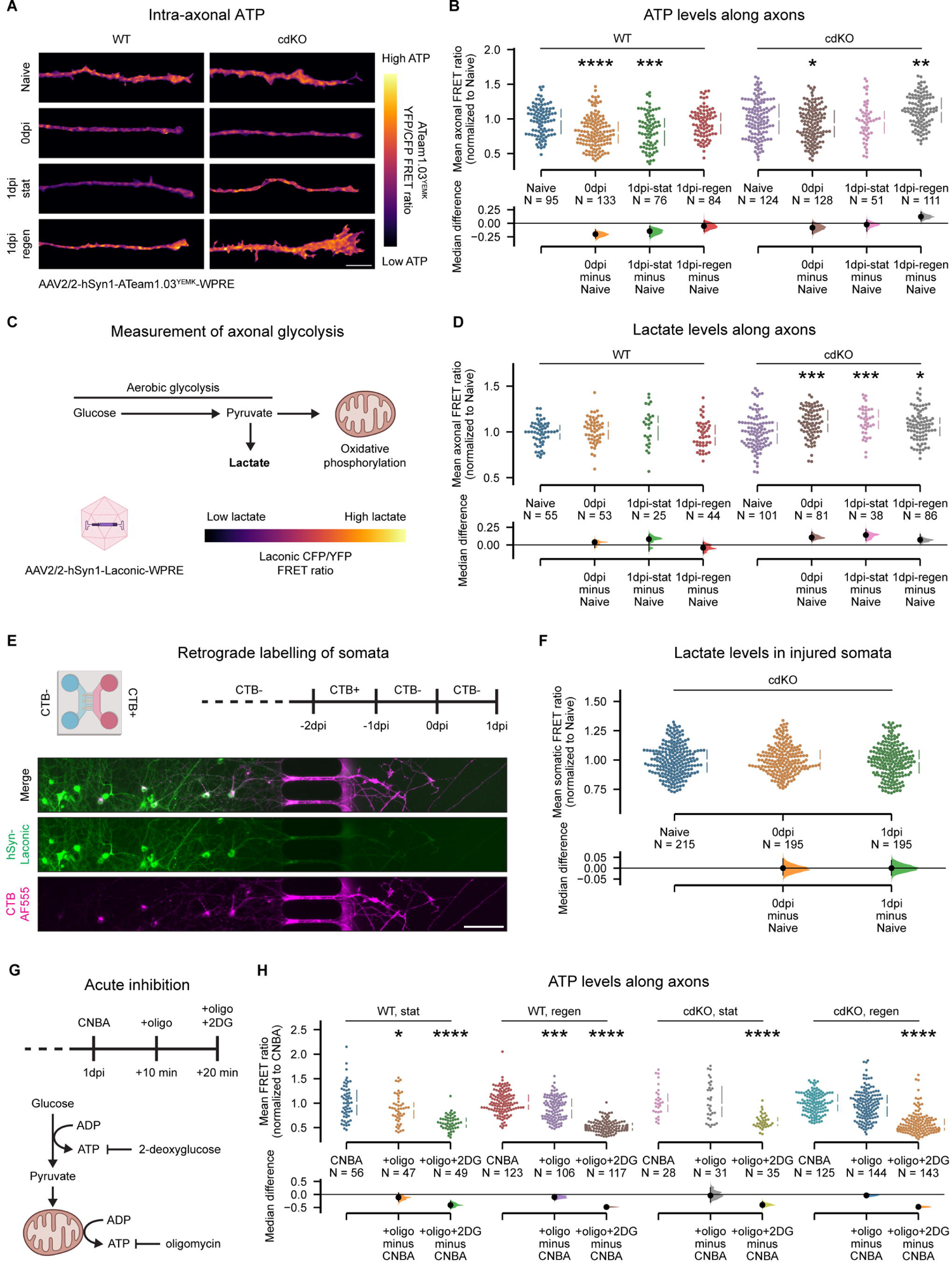
Regeneration of cdKO axons is accompanied by increased ATP concentration, supported by axonal glycolysis. (A) Representative images of axonal endings of WT and cdKO neurons, depicted as pseudo-colour representations of the FRET ratio of the ATP biosensor ATeam1.03^YEMK^. Scale bar 10 µm (B) Quantification of the FRET ratio along axon shows restoration of ATP only in regenerating axons for WT neurons and in both static and regenerating axons for cdKO neurons. (C) Schematic representation of glucose metabolism, resulting in lactate production via aerobic glycolysis (Warburg effect) or pyruvate import into mitochondria for oxidative phosphorylation. Axonal lactate is measured with the biosensor Laconic. (D) Quantification of axonal intracellular lactate concentration reveals upregulated aerobic glycolysis in cdKO axons following injury and at 1 dpi, regardless of regeneration outcome. (E) Schematic representation of retrograde labelling of microgroove-crossing neurons with Cholera Toxin B (CTB), allowing to specifically label the somata of the RGCs that will be injured. Scale bar 100 µm (F) Quantification of intracellular lactate within the somata of cdKO injured RGCs shows no upregulation of aerobic glycolysis in the somatic compartment after injury and at 1dpi. (G) Schematic representation of the sequential inhibition of oxidative phosphorylation and glycolysis, via administration of 10 µM oligomycin and of 50 mM 2-deoxyglucose, respectively, in the axonal compartment. (H) Quantification of the FRET ratio along axons in normal complete Neurobasal-A medium (CNBA) and in presence of oligomycin (+oligo) or oligomycin and 2-deoxyglucose (+oligo+2DG) for each axon class. A significant decrease in ATP concentration is detected in WT axons after inhibition of oxidative phosphorylation, but not in cdKO axons. Data from 5 (B) or 4 (D, H, F) independent experiments, presented as median ± 25-75^th^CI and bootstrap 95% CI versus relative naive (A, D, F) or relative CNBA (H). Kruskal-Wallis ANOVA. *p<0.05, **p<0.01, ***p<0.001, ****p<0.0001. See also Fig S3.

Considering that both static and regenerating cdKO axons showed higher ATP at 1dpi as compared to WT (Fig S3B), irrespective of regeneration outcome and mitochondrial phenotype, we hypothesized that the deletion of *Pten* and *Socs3* leads to an increase in non-mitochondrial energy production within the axons. Under the Warburg effect^20,21^, aerobic glycolysis produces 2 ATP molecules in the cytosol during the conversion of glucose into lactate. While less efficient than oxidative phosphorylation, it has a faster turnover ^20^. To assess whether mitochondria-independent energy production takes place in cdKO axons, we expressed the biosensor Laconic (*AAV2/2-hSyn1-Laconic-WPRE-hGHp*) and measured the axonal concentration of lactate (Fig 3C). Uninjured axons showed a comparable concentration of lactate in WT and cdKO neurons (Fig S3C). We did not detect any significant change in axonal lactate in WT axons at 0dpi and 1dpi as compared to naive. On the contrary, cdKO axons significantly upregulated lactate production after injury at 0dpi. This increase was maintained at 1dpi in static axons and, strikingly, also in cdKO regenerating axons (Fig 3D). In these, the concentration of lactate was higher than that in regenerating WT axons (Fig S3D).

Next, to determine whether the upregulation of glycolysis in cdKO neurons is a local compensatory effect within the axons rather than a neuron-wide phenotype, we measured the concentration of lactate within the soma of the injured RGCs. To achieve this, prior to injury, we retrogradely traced the RGCs of which the axons extend into the axonal channel and thus that will be injured during axotomy (Fig 3E). Strikingly, no somatic upregulation of lactate production was detected in injured RGCs at both 0dpi and 1dpi as compared to naive (Fig 3F), suggesting that the upregulation of aerobic glycolysis is a local axon-restricted effect.

Finally, to determine the relative contribution of oxidative phosphorylation and glycolysis to the pool of axonal ATP at 1dpi, we sequentially inhibited both processes in the axonal compartment with oligomycin (oligo) and 2-deoxyglucose (2DG), respectively (Fig 3G). In WT axons we detected a significant reduction in ATP after inhibition of oxidative phosphorylation and a further reduction upon additional inhibition of glycolysis (Fig 3H). On the other hand, in dKO axons we did not observe a significant decrease in ATP with oligomycin, but rather only after combined inhibition of glycolysis (Fig 3H). This suggests that cdKO axons can sustain ATP via glycolysis alone. The relative percentages of ATP production calculated from this experiment are depicted in supplement (Fig S3E). In summary, deletion of *Pten* and *Socs3* improves the maintenance of axonal ATP production after injury and during initiation of regrowth. This enhancement is provided by a local upregulation of glycolysis, which provides the bulk of the ATP.

### Downregulation of axonal glycolysis reverses the regeneration phenotype upon

#### *Pten* and *Socs3* deletion

To address whether the increased glycolysis upon injury in cdKO axons is necessary for regeneration, we downregulated the glycolytic flux in the axonal compartment following axotomy. This was achieved via the substitution of glucose for galactose in the axonal medium from 0dpi until 1dpi (Fig 4A). Galactose is an alternative sugar, which can be converted into glucose-6-phosphate (G6P) through the Leloir pathway to feed into glycolysis^22–24^. In the absence of glucose, the conversion of galactose into G6P becomes rate-limiting, reducing the flux of glycolysis and reducing glycolytic ATP^24^ (Fig 4A). With galactose as a substrate, cells have been shown to rely more heavily on mitochondrial oxidative phosphorylation to meet energy demands^24,25^.

**Figure 4:**
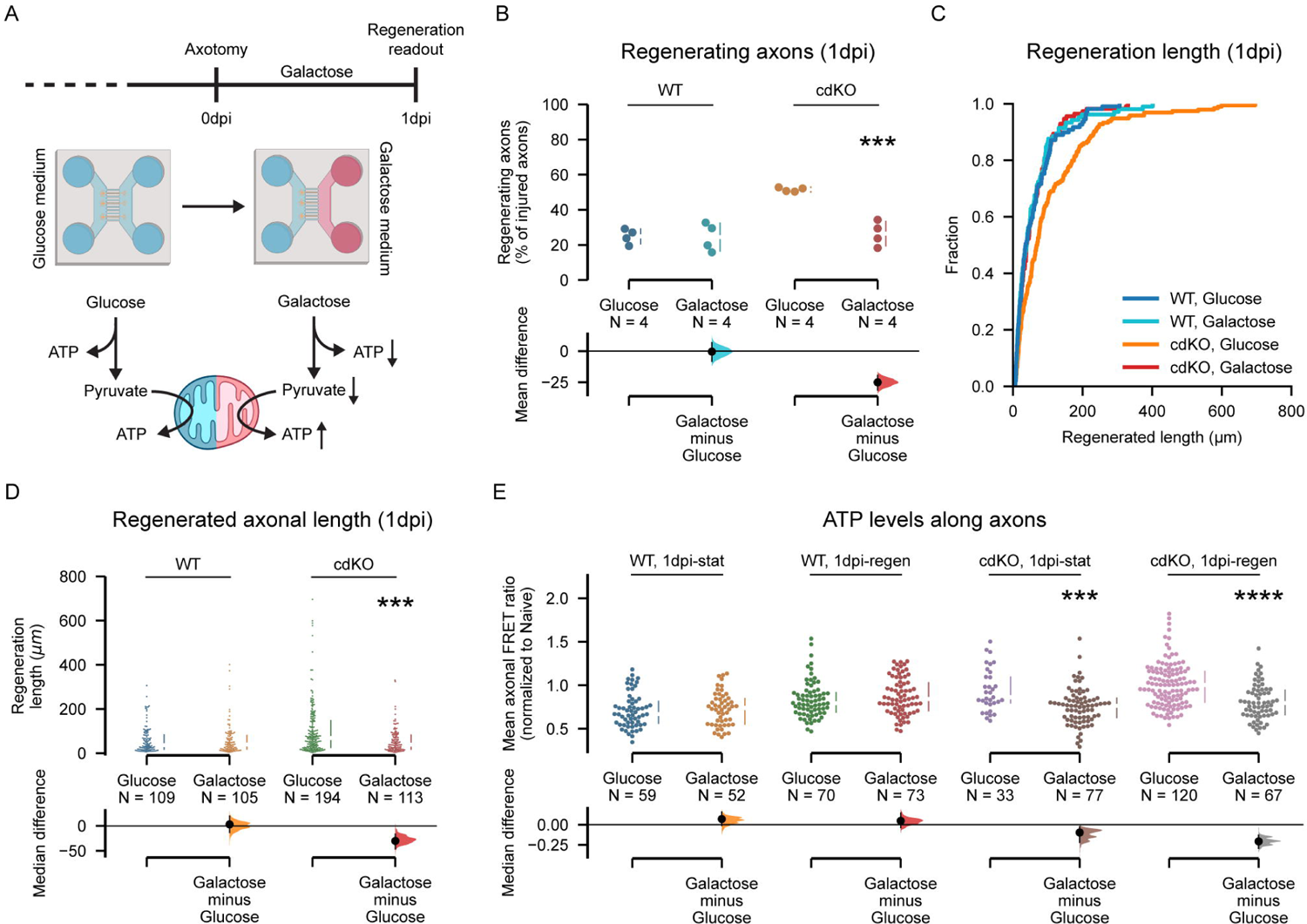
Impairment of axonal glycolysis abolishes the regeneration phenotype of cdKO axons, via decreased axonal ATP. (A) Schematic representation of the axonal glycolysis downregulation. Glucose in the medium is switched to galactose after axotomy and maintained until 1dpi. Galactose is processed via glycolysis at a slower rate, leading to reduced glycolytic ATP and a relative induction of mitochondrial ATP. (B) Quantification of the percentage of regenerating axons shows that galactose treatment does not affect WT axons but reverses the enhanced regeneration phenotype of cdKO axons. (C) Cumulative distribution of axonal lengths reveals that galactose treatment reduces the regrowth length in cdKO axons at 1dpi. (D) Quantification of the regenerated axonal length shows that galactose treatment induces a significant decrease in the average axonal regenerated length in cdKO axons at 1dpi. (E) Quantification of the FRET ratio along axons reveals that downregulation of axonal glycolysis does not impact the ATP concentration of WT axons at 1dpi but reduces the ATP concentration of both static and regenerating cdKO axons. Data from 4 independent experiments, presented as mean ± SD (B) or median ± 25-75^th^CI (D, E) and bootstrap 95% CI versus glucose. Student t-test (B), Mann-Whitney U-test (D, E). ***p<0.001, ****p<0.0001. See also Fig S4.

Galactose treatment did not alter the limited regeneration phenotype of WT axons, both in terms of percentage of regenerating axons (Fig 4B) and as well as length (Fig 4C-D). On the other hand, the downregulation of axonal glycolysis in cdKO axons completely reversed the induced regeneration phenotype of *Pten* and *Socs3* co-deletion. Both the percentage of regenerating cdKO axons as well as their length significantly decreased to levels comparable to WT axons when grown in galactose medium (Fig 4B-D). Also, the percentage of degenerating and static cdKO axons decreased to levels comparable to WT when in galactose medium (Fig S4A-B).

To determine whether the downregulation of glycolysis impacted the intra-axonal energy levels, we measured the concentration of ATP during galactose treatment. Limiting glycolysis did not impact ATP production in both static and regenerating WT axons at 1dpi (Fig 4E). On the other hand, both static and regenerating cdKO axons showed significantly lower ATP levels at 1dpi in galactose medium as compared to glucose (Fig 4E). Finally, we labelled mitochondria with MitoTracker to assess whether the treatment affected the mitochondrial phenotype. Notably, galactose did not alter mitochondrial transport and density in cdKO axons (Fig S4C,E). At 1dpi, we measured comparable mitochondrial trafficking between regenerating axons in the glucose and galactose condition (Fig S4C). Only a minor increase in mitochondrial length was measured in regenerating axons when grown in galactose medium (Fig S4D). This is likely due to a compensatory increase in mitochondrial fusion to sustain oxidative phosphorylation. In summary, these findings support the hypothesis that local upregulation of axonal glycolysis underlies the enhanced regrowth of axons upon deletion of *Pten* and *Socs3*.

## DISCUSSION

Axonal regeneration is an energy demanding process that has been reported to rely on the transport of healthy mitochondria to the axons^6,7,10,26^. In this study, we have shown that axotomy-induced axonal regeneration of cultured RGCs is characterized by the restoration of mitochondrial axonal transport and that the increased regeneration upon *Pten* and *Socs3* deletion is accompanied by an upregulated mitochondrial trafficking. While WT axons rely on these mitochondria to meet energy demands, we demonstrated that upon deletion of *Pten* and *Socs3*, local glycolysis in the axons significantly supports energy production. Limiting the glycolytic flux with galactose medium impaired their energy production and completely reversed the regeneration phenotype of *Pten* and *Socs3* co-deleted axons.

### Circumventing an irreversible energy crisis

Injury has been reported to alter the axonal environment, leading to mitochondrial depolarization and dysfunction^7,10^. This causes a depletion of axonal ATP, which is associated with Wallerian degeneration^27,28^. According to the established paradigm of axonal regeneration, mitochondria accumulate at the growth cone and provide energy to fuel and support axonal elongation. Nonetheless, any delay or impairment in the mobilization of mitochondria and their transport into the axons can lead to an irreversible energy crisis, preventing the initiation of regeneration or, even worse, causing degeneration^27,28^. Unfortunately, mitochondrial transport is progressively decreased during neuronal maturation, with increased mitochondrial docking at the site of synapses, mediated by the adaptor proteins connecting the mitochondria to the kinesin and dynein motors, such as Miro and syntaphilin^10,13,26^. This has been ascribed as one of the mechanisms restricting axonal regeneration in the mammalian CNS^1,2^.

In accordance, in our setup of cultured RGCs, we have shown that restoration of axonal mitochondrial transport and morphology is indeed required for axotomy-induced axonal regrowth. Unlike a previous report by Cartoni et al. on embryonic cortical neurons^6^, we did not detect a higher basal mitochondrial transport in uninjured axons upon *Pten* and *Socs3* co-deletion, but only in regenerating ones. This might be due to the higher degree of maturity and mitochondrial docking of the neurons in our setup or a difference between the neuronal classes. Strikingly, the enhanced regeneration associated with *Pten* and *Socs3* deletion is accompanied by an immediate upregulation of axonal glycolysis, which provides a considerable and essential fraction of the energy. The inability to upregulate glycolysis and combine it with the restoration of mitochondrial function, impairs the enhanced regeneration otherwise observed in cdKO axons.

### Glycolysis: an essential gear in the axonal machinery

Neurons predominantly rely on mitochondrial metabolism to meet energy demands. Indeed, differentiation of neuronal progenitor cells into neurons is accompanied by a shift from glycolysis to oxidative phosphorylation^29,30^. Metabolic support from glycolytic glial cells, i.e., astrocytes and oligodendrocytes, involves lactate production that is taken up by neurons, converted to pyruvate, and used for oxidative phosphorylation^31,32^. Lately, researchers started to challenge this notion, showing that glucose uptake and glycolysis are essential for basal neuronal functions^23,33^. Recent evidence in *C. elegans* suggests that neurons autonomously carry out glycolysis and can regulate it at a subcellular compartment level^34^. Neuronal stimulation has been reported to induce glycolysis rather than oxidative phosphorylation *in vivo* in mice, temporarily making neurons net exporters of lactate^35^. Moreover, fast axonal transport has been described to be fuelled by the glycolytic metabolon assembled on the vesicles rather than by mitochondria^36,37^, and this requires the reduction of pyruvate into lactate^38^. Likewise, both developmental axonal growth and retraction *in vitro* have been shown to rely on glycolysis, in embryonic chick dorsal root ganglia (DRG) and embryonic cortical rat neurons, respectively^39,40^. Because of the short half-life of ATP and its low coefficient of diffusion^41^, it appears that high energy-consuming processes, like cytoskeletal remodelling and kinesin-mediated movement in the axon, require ATP produced in extreme proximity. Therefore, tight coupling of the energy-draining machinery with the enzymes of the ATP-producing steps of glycolysis can provide local cytosolic ATP^39,40^. Taken together, this might explain why increased glycolytic ATP in our cdKO axons is beneficial after axonal injury, to remodel the cytoskeleton during growth-cone formation and to fuel vesicular transport of somatic material into the axon.

### A strengthened glycolysis-mitochondria axis

In the context of axonal injury, the ability to upregulate glycolysis might thus fill the energy gap caused by depolarized and dysfunctional mitochondria, counteracting degeneration, and initiating regrowth. A recent study reported that glycolysis is activated in the peripheral nervous system (PNS) after injury ^42^, not only in Schwann cells^43^, but also in axons and that inhibition of glycolysis increased Wallerian degeneration^42^. These data support our findings in regeneration-prone CNS neurons, showing injury-induced glycolytic upregulation in *Pten* and *Socs3* deleted RGCs.

In this scenario of upregulated glycolysis, mitochondria are not redundant, as they exert a wide array of cellular functions besides producing ATP. These include calcium buffering, redox homeostasis, and the generation of precursors for the synthesis of macromolecules essential for cellular survival^44,45^. While the latter have not been studied as well in neurons as in other cell types, it has been shown that neuronal lipid synthesis requires cooperation between mitochondria and the endoplasmic reticulum^45,46^. In proliferating cells, under the Warburg effect, aerobic glycolysis provides most of the cellular ATP^20,21^. Nonetheless, mitochondria are still functional and via the Krebs cycle have the critical role to provide acetyl-CoA for fatty acid synthesis and aspartate for amino acid synthesis. Neurons could benefit from this “separation of tasks” during axonal elongation, with upregulated glucose metabolism to generate ATP and reprogramming of mitochondrial metabolism towards anabolic tasks. As such, the integration of a high glycolytic flux with the maintenance of healthy mitochondria, as seen in regenerating cdKO axons, could provide an ideal biochemical environment for regrowth.

### Limitations of the study

Our study was conducted in an *in vitro* setting, which can cause adaptations to the basal neuronal metabolism as compared to *in vivo*. While we employed mixed cultures containing both retinal neurons and glia, the latter were absent from the axonal compartment, preventing possible axon-glia metabolic interactions that could occur *in vivo*. Indeed, Schwann cells have been shown to support regeneration in the PNS via increased glycolytic activity and lactate shuttle to the axons^43^. On the other hand, in the context of optic nerve crush, resident oligodendrocytes were shown to undergo demyelination and partial cell death^47^. How this disrupted environment affects the metabolic support to injured and regrowing axons is unclear, but at least during the first phases of regeneration it is reasonable to speculate that the axons could rely primarily on intrinsic glucose metabolism rather than metabolic coupling with glia.

Culture media optimized for neuronal growth, such as Neurobasal-A, often contain concentrations of glucose which are hyper-physiological. Not much is known about how this affects neuronal metabolism, but the use of high-glucose media has been shown to induce a lower reliance on mitochondrial metabolism in yeast and mammalian cell lines, in what is known as the Crabtree effect^48^. Nonetheless, despite the relatively high levels of basal glycolysis, we detected an increase in lactate production and glycolytic flux after injury specifically in axons rather than somas and in *Pten* and *Socs3* co-deleted neurons rather than WT ones. Whether the magnitude of this upregulation would be similar *in vivo* is unclear and further research is required to elucidate this, but the evidence presented in this study supports the hypothesis of this being a specific injury-induced mechanism underlying axonal regeneration.

### Impact and future perspectives

While the deletion of *Pten* and *Socs3* affects a wide array of cellular functions and is not suited for clinical translation, the targeting of glycolysis could support and be synergistic with other less invasive strategies to induce axonal regrowth, such as targeting of IL6 signalling or the overexpression of protrudin^49,50^. In summary, the manipulation of glycolysis and of axonal metabolism at large, could provide new therapeutic avenues to unlock the regeneration potential of the mammalian CNS.

## ACKNOWLEDGMENTS

We thank Evelien Herinckx and Arnold Van Den Eynde for the animal caretaking, Iene Kemps and Marijke Christiaens for technical support and Joana Santos, Cristiano Lucci, Romana Fato and Christian Bergamini for the critical discussions. We thank Dr. Xandra Pereiro and Prof. Dr. Elena Vecino for their support and advice during the initial setup of the culture protocol. Lu.M, S.B, A.V.D and L.D.G. were supported by personal fellowships funded by the Research Foundation Flanders (FWO, Belgium) (fellowships 1S42720N, 1165020N, 1S94218N, 12I3820N). This research was funded by the FWO (project G082221N) and the KU Leuven Research Council (C14/22/074).

## AUTHOR CONTRIBUTIONS

Conceptualization: LuM, SB, AVD, LDG, KF and LM; Methodology: LuM; Investigation: LuM; Data curation: LuM, SB, LM; Formal analysis: LuM; Visualization: LuM; Software: LuM; Writing—original draft preparation: LuM; Writing—review and editing: SB, AVD, LDG, KF, LM; Funding acquisitions: LuM, SB, AVD, LDG, KF and LM; Visualization: LuM; Project administration: LuM, LM: and Supervision: LDG, KF, LM.

## STAR METHODS

### KEY RESOURCE TABLE

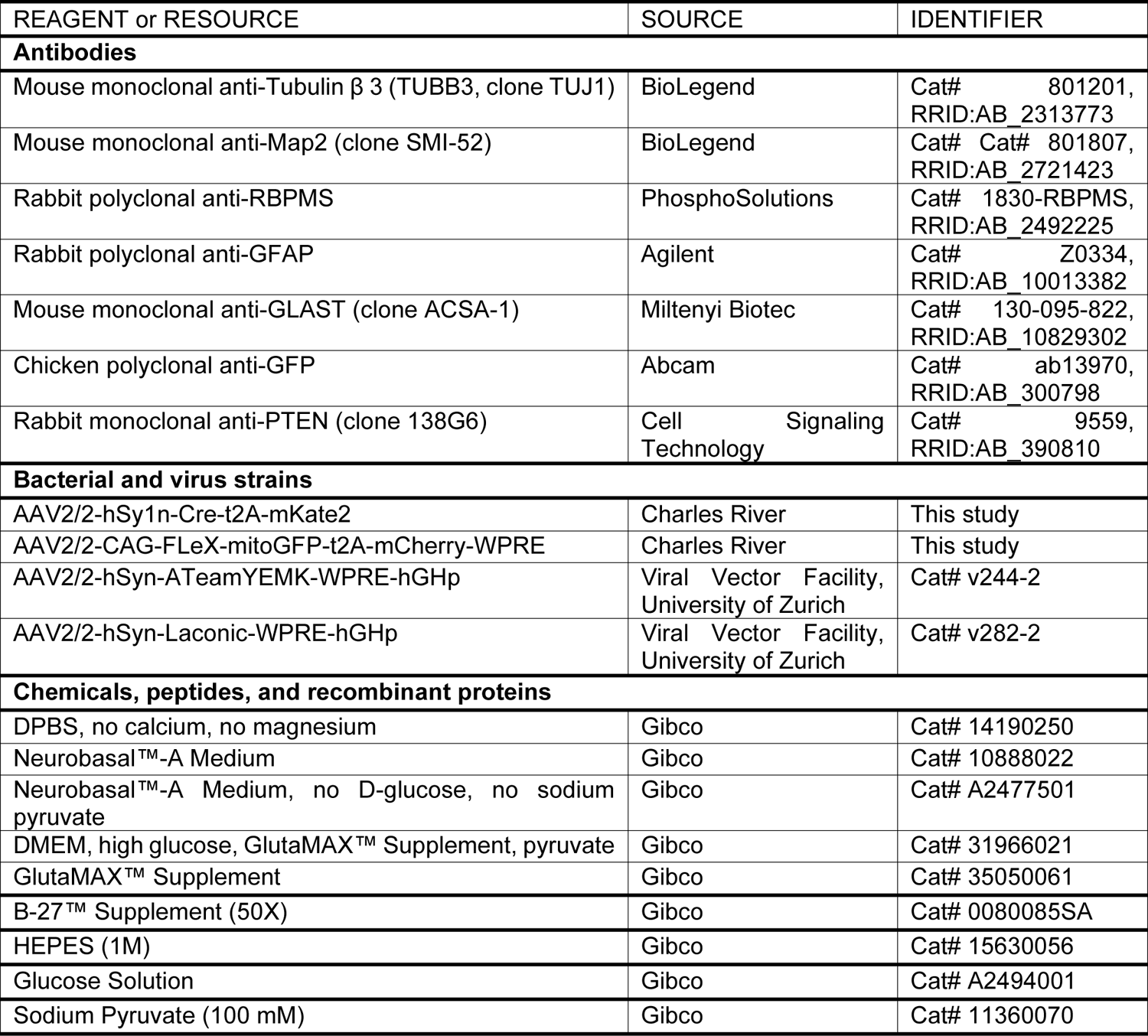
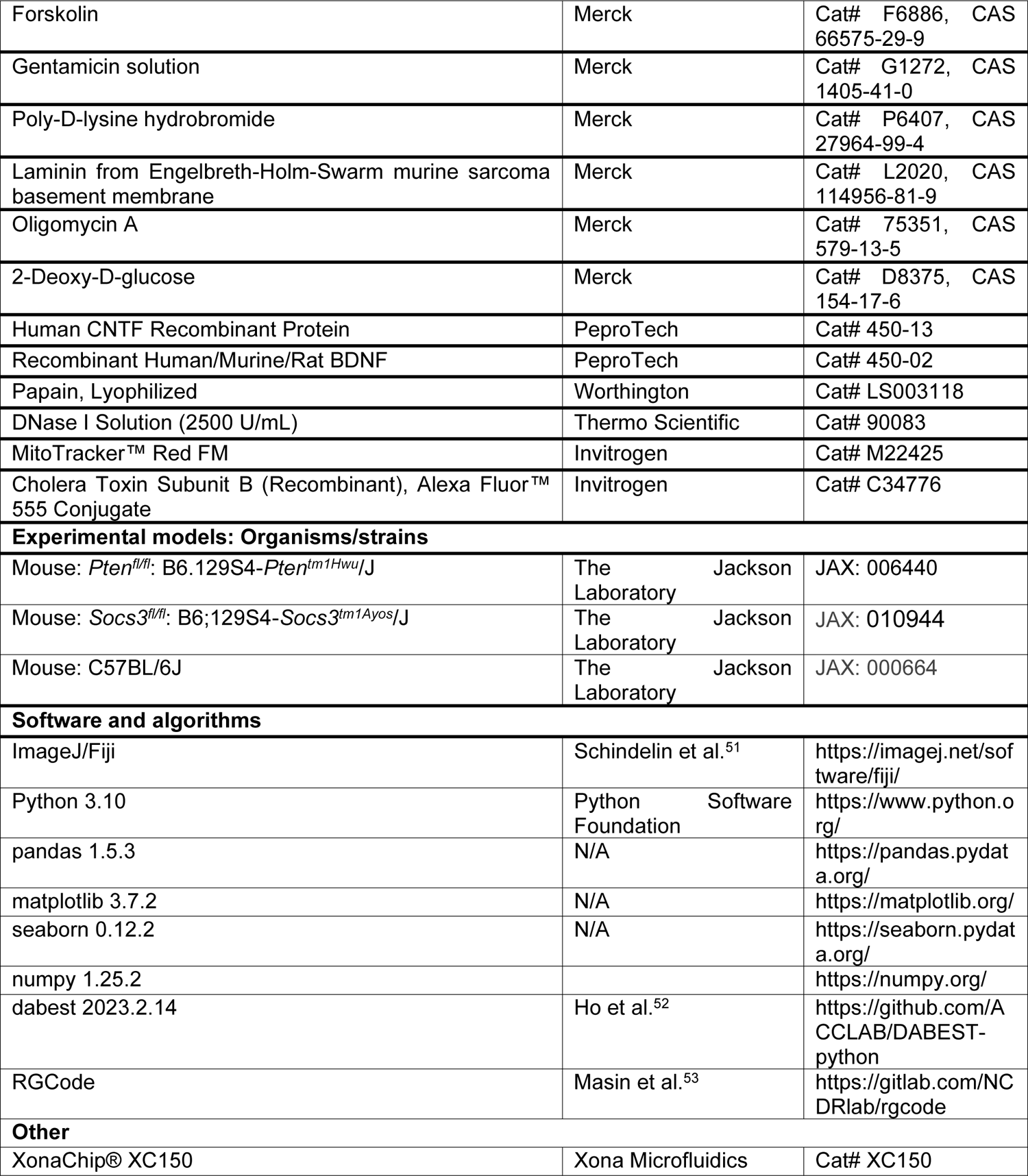

### RESOURCE AVAILABILITY

#### Lead contact

Further information and requests for resources and reagents should be directed to and will be fulfilled by the lead contact, Lieve Moons

#### Materials availability

All non-commercial reagents or mouse lines used in this paper are available from the lead contact upon request.

#### Data and code availability

- All original data reported in this paper will be shared by the lead contact upon request.
- This paper does not report original code.
- Any additional information required to reanalyze the data reported in this paper is available from the lead contact upon request.

### EXPERIMENTAL MODEL AND STUDY PARTICIPANT DETAILS

#### Animals and housing

Experiments were performed on male and female postnatal day (P) 2-3 *Pten^fl/fl^;Socs3^fl/fl^* mice, obtained by crossing *Pten^fl/fl^*and *Socs3^fl/fl^* mice^54,55^, which were purchased from Jackson Laboratories. For mitochondrial trafficking experiments, male and female C57Bl6 mice were used to generate wild-type (WT) cells (see viral vector section of the Methods).

Mice were housed in a temperature-, light- and humidity-controlled environment, under a 12h light/dark cycle and with *ad libitum* access to food and water. All animal experiments were approved by the Institutional Ethical Committees for Animal Experimentation of KU Leuven and conducted following strictly the European and Belgian legislations.

#### Primary retinal cultures

##### Retinal cell isolation

Retinal cells were isolated from P2-P3 mouse pups following previously reported methods with minor modifications^56^. Retinae were dissected from the eyes in warm sterile Dulbecco PBS (DPBS) and collected during the procedure in warm Dulbecco’s Modified Eagle Medium (DMEM) supplemented with 50 µg/mL gentamycin (gDMEM). After dissection, retinae were dissociated with 16 U/mL of papain in 1 mL of gDMEM for 30 minutes, gently triturated with a pipet tip, filtered through a 40 µm cell strainer and resuspended in 200µL of complete Neurobasal-A medium (CNBA) (Neurobasal-A, 2 mM GlutaMAX, 10 mM HEPES, 2% B27, 5 µM forskolin, 50 µg/mL gentamicin) with the addition of 45 U/mL of DNAse to prevent clumping.

##### Microfluidic seeding and culture

Retinal cells were seeded on XONA XC150 microfluidic devices, which were previously sequentially coated with poly-D-lysin (0.1 mg/mL in water) and laminin (2 mg/mL in DPBS), both overnight at room temperature. Briefly, 10 µL of medium containing 250.000 cells was added to the somatic channel of microfluidic device (MFD). Cells were allowed to attach for 1 hour and subsequently the wells of the somatic chamber of the MFD were filled with somatic CNBA (sCNBA: CNBA, 2 ng/mL CNTF, 5 ng/mL BDNF) and on the axonal chamber with axonal CNBA (aCNBA: CNBA, 20 ng/mL CNTF, 50 ng/mL BDNF). A small fluidic gradient was kept across the grooves during normal culture by adding 150 µL of medium per well on the somatic side and 120 µL per well on the axonal one. In each well, half of the medium was replaced with fresh one every other day during the extent of the culture. Cells were kept in a humidified incubator supplied with 5% CO_2_ at 37°C.

##### Viral constructs and transduction

Retinal cells were transduced at day in vitro 1 (DIV1) with AAV vectors diluted in sCNBA and added on the somatic chamber only. For the imaging of ATP levels, cells were transduced with an *AAV2/2-hSyn1-ATeam^YEMK^-WPRE-hGHp* vector (1.0×10^10^ GC/mL final titre in medium) and for the imaging of lactate levels with a *AAV2/2-hSyn1-Laconic-WPRE-hGHp* vector (6.7×10^9^ GC/mL final titre in medium). Only for the cdKO condition, an *AAV2/2-hSyn1-Cre-t2A-mKate2* vector (1.8×10^9^ GC/mL final titre in medium) vector was used to in combination with the reporters to induce gene recombination and deletion in *Pten^fl/fl^;Socs3^fl/fl^* cells. To evaluate knockout efficiency and axonal regeneration in Fig 1 and Fig S1, ATeam was used as a fluorescent reporter. In this case, only the YFP fluorophore was excited and is thus reported as hSyn-YFP in the figures and text. The use of a hSyn promoter ensures a high degree of RGC specificity.

For mitochondrial imaging, cells were transduced with a Cre-dependent *AAV2/2-CAG-FLeX-mitoGFP-t2A-mCherry-WPRE* vector (2.7×10^10^ GC/mL final titre in medium). This vector encodes a cytosolic mCherry to label the entire neuron and a tagged GFP to label mitochondria. Mitochondrial targeting of GFP is achieved by fusing the target sequence of *Cox8a* on the N-terminus of GFP. Since this vector is Cre-dependent, to avoid recombination of *Pten* and *Socs3* in WT cells, C57Bl6 cells were used instead of *Pten^fl/fl^;Socs3^fl/fl^*ones in all experiments depicted in Fig 2 and Fig S2 for the WT condition.

## METHOD DETAILS

### *In vitro* procedures

#### Axonal injury

Axonal injury was performed between DIV12 and DIV14, via vacuum aspiration of the axonal medium. Briefly, a thin gel loading tip was attached to an aquarium pump and used to empty the well of the axonal chamber. After, the tip is moved to the entrance of the axonal channel to remove its medium. The operation is performed twice, once per channel entrance. Immediately after, the axonal chamber is refilled with fresh medium. Vacuum aspiration severed the axons in the axonal channel at approximately 100 µm from the exit of the grooves. In all experiments, at 0dpi and 1dpi, axons were analysed in the space between the groove exit and their most distal tip in the axonal channel.

#### Retrograde tracing

Retrograde tracing was performed by supplementing the axonal medium with Alexa-conjugated Cholera Toxin B (CTB-AF555) to a final concentration of 0.1 mg/mL in aCNBA, 48h before injury. During CTB exposure, the fluidic grading was increased to 150µL per well in the somatic chamber and 80 µL per well on the axonal one, to prevent flow of CTB to the axonal side. The CTB-supplemented medium was completely washed and replaced with aCNBA the day after, 24h before axonal injury.

#### Metabolic pathway inhibition

To determine the relative fractions of ATP production at 1dpi, we measured the axonal ATP concentration (see below) sequentially in axonal medium (aCNBA), in aCNBA supplemented with 10 µM of oligomycin and in aCNBA supplemented with 10 µM of oligomycin and 50 mM of 2-deoxyglucose. To locally reduce glycolysis in the axons, right after axotomy we replaced aCNBA with galactose medium (gCNBA: glucose- and pyruvate-free Neurobasal-A, 0.27 mM sodium pyruvate, 2 mM GlutaMAX, 10 mM HEPES, 2% B27, 5 µM forskolin, 50 µg/mL gentamicin, 20 ng/mL CNTF, 50 ng/mL BDNF). During treatment, the small fluidic gradient was reversed to prevent flow of glucose medium from the somatic chamber to the axonal one.

#### Mitochondrial labelling

Due to spectral incompatibility, for the experiments concerning galactose treatment, when combining the imaging of ATP levels with mitochondrial transport, mitochondria were labelled with MitoTracker Red FM instead of mitoGFP. On the day of imaging, cells were incubated with 100 nM MitoTracker for 45 minutes in medium in both microfluidic compartments and washed prior to imaging.

#### Immunocytochemistry

For immunostaining, cells were fixed for 10’ with 4% paraformaldehyde in PBS (137 mM NaCl, 2.7 mM KCl, 10 mM Na_2_HPO_4_, 1.8 mM KH_2_PO_4_ in distilled water, pH 7.4). All stainings were carried out within the MFDs. Cells were permeabilized with 0.1% Triton-X100 in PBS (PBST), thrice for 5’. Thereafter, antigen blocking was performed with 10% pre-immune donkey serum (PID) in PBST for 45’. Following, primary antibodies were diluted to the desired concentrations in PBST, supplemented with 10% PID and incubated overnight at room temperature. For labelling of PTEN, the incubation was carried out for 4 days at 4°C with the addition of 10% PID and 0.04% NaN_3_. After primary antibody binding, cells were incubated for 2 hours at room temperature with Alexa-conjugated secondary antibodies (1:200 in PBST). Map2 was labelled using an Alexa647 pre-conjugate primary antibody incubated overnight at 4°C after the secondary antibodies were bound, to prevent cross-reactivity with primary antibodies of the same host. After final washing, the cells were incubated for 30’ with DAPI (1:1000 in PBS) to label the nuclei. Samples were stored in PBS until imaging, which was performed within 24h from the staining.

#### Imaging

All live imaging was performed on a Zeiss LSM900 with Airyscan 2, with a STXG incubation system (Tokai Hit) maintaining the samples at 5% CO_2_ and 37°C.

#### Epifluorescence imaging

The entire MFD, as in somatic channel, microgrooves and axonal channel, was imaged before, after injury and at 1dpi. Imaging was performed in epifluorescence mode with a Plan-Apochromat 20x/0.8 M27 objective to minimize imaging time and phototoxicity. This way, the whole MFD was imaged in the span of 5 minutes.

#### Confocal imaging

All confocal imaging was performed with a Plan-Apochromat 20x/0.8 M27 objective in CO-2Y mode. Mitochondrial trafficking was imaged via timelapse, carried out for 10’ per frame at a frequency of 0.5Hz. ATP and lactate biosensor imaging was performed with 405 nm excitation and with emission filters set manually to 450-512 nm for CFP and 512-573 nm for YFP. For Tuj1 and Map2 labelling only, MFD were imaged using an Olympus FV1000 confocal microscope, with a UPLSAPO 20X/0.75 objective.

### Image analysis

#### Axonal regeneration

Whole-MFD images of the somatic and axonal compartments at 0dpi and 1dpi were aligned via descriptor-based image registration with ImageJ. On these overlaid images, comparing 0dpi and 1dpi, axons were counted and scored manually in their classes, being regenerating, degenerating or static. The length of regenerating axons was measured by manually tracing them with the ImageJ plugin NeuronJ, from the most distal tip at 1dpi until the site of injury (i.e. the distal tip at 0dpi).

#### Mitochondrial kymographic analysis

Timelapses were stabilized to remove stage drift via descriptor-based image registration with ImageJ. Kymographs were generated and analysed manually using the Kymolyzer set of macros in ImageJ^57^. Mitochondrial length and occupancy were measured with an ImageJ script. The axon trace from the kymograph analysis was loaded and the axonal mitochondria were segmented via Otsu thresholding. After, the mitochondrial morphological statistics were calculated via the Analyze Particles function. The major axis of the ellipse fit on the segmented particle was used as a measurement of mitochondrial length. Averages of mitochondrial trafficking and length statistics per axon are calculated in Python with the *pandas* library. As previously reported^17^, the mitochondrial axonal occupancy was defined as the fraction of axonal length occupied by mitochondria. Occupancy was calculated as the sum of the length of all mitochondria per axon divided by the axonal length.

#### ATP and lactate analysis

For the analysis of ATP and lactate concentration, the ratios of YFP/CFP and CFP/YFP, respectively, were calculated in Python with the numpy and scikit-image libraries after application of a median filter to reduce noise. For quantification, regions of interest (ROI) were manually drawn and the average ratio intensity per axon or soma was measured in ImageJ. To minimize inter-experiment scattering of the datapoints, per independent experiment, datapoints were normalized against the median of their relative naive group. Only for representative images in the figure panels, for ease of visualization and comparison of the pseudo-colour, the non-specific FRET signal outside the axons was removed by Otsu thresholding of the YFP signal and using the “Clear Outside” function of ImageJ.

#### Quantification of knock-out efficiency

To quantify the efficiency of AAV-mediated Cre knockout of *Pten* and *Socs3*, the somata of the RGCs were segmented using RGCode, a deep learning tool developed in-house^53^. For this purpose, a new segmentation model was trained from manual annotations on the dataset. Segmentation was run on images of the entire somatic channel of the MFD. After segmentation, the mean fluorescence intensity of PTEN was measured per RGC soma. A threshold of mean somatic intensity was manually defined per independent experiment to classify RGCs as PTEN+ or PTEN- and calculate their relative percentage.

## QUANTIFICATION AND STATISTICAL ANALYSIS

Detailed information on the number of samples and axons used are reported in the legends of the figures. The number of *N* axons per condition/timepoint is reported in the graph under each group. These are accumulated several independent experiments, of which the specific number is reported in the figure legends. In experiments concerning percentages of axons per microfluidic device, *N* corresponds to the number of independent experiments rather than axons. All data analysis was performed on raw micrographs, which were not saturated during acquisition. For visualization purposes in figure panels, some images were inverted and/or contrast-enhanced by reducing the white point (or increasing the black point when inverted). In this case, the same magnitude of enhancement was applied to all shown images when compared. Both statistical tests such as ANOVA, t-test and U-test, as well as bootstrapping were used for significance testing and the related details are reported in the figure legends. The median was used together with Kruskal-Wallis ANOVA if any of the groups in the experiment didn’t pass the Shapiro-Wilk normality test. Otherwise, the mean and Welch ANOVA were used. As the short recording time leads to zero-inflated data for technical reasons during the measurement of mitochondrial moving frequency, the mean and Welch ANOVA were used despite non-normal data. All numerical data processing, plotting, statistical tests and bootstrapping were conducted in Python with the libraries *pandas*, *seaborn*, *matplotlib*, *scipy* and *dabest*^58^. A p value of <0.05 was considered significant.

## SUPPLEMENTAL INFORMATION TITLE AND LEGENDS

**Figure S1:** Validation of the retinal cell culture and microfluidic model. (A) Immunocytochemistry for Tuj1 and RBPMS at 14 days *in vitro* (DIV) identifies retinal ganglion cells (RGCs), co-labelled by both markers, in the mixed retinal cell culture. Scale bar 25 µm. (B) Immunocytochemistry for Tuj1 and Map2 at 14DIV identifies Tuj1+ RGCs in the somatic compartment and their axons in the axonal one. Importantly, the dendritic marker Map2 is restricted to the somatic compartment. The edges of the microgrooves are delineated by the orange dotted line. Scale bar 100 µm. (C) Immunocytochemistry for GFAP and GLAST at 14DIV revels the presence of retinal macroglia, co-labelled by both markers. Scale bar 25 µm (D) Immunolabelling for PTEN and Tuj1 shows specific loss of PTEN within Tuj1+/hSyn-YFP+ RGCs upon transduction with an AAV2/2-Syn-Cre viral vector (cdKO) as compared to WT cells, indicating specific recombination and gene knock-out. hSyn-YFP is encoded by *AAV2/2-hSyn1-ATeam^YEMK^-WPRE-hGHp*, see Methods. Scale bar 50 µm. (E) Quantification of PTEN signal in the somata of hSyn-YFP+ RGCs showes that on average 98.7% of the RGCs are positive for PTEN at 14DIV in the WT condition as compared to only 2% in the cdKO culture, indicating that an efficient knock-out is achieved. (hSyn-YFP encoded by *AAV2/2-hSyn1-ATeam^YEMK^-WPRE-hGHp*) (F) Graphical representation of the percentages of regenerating (green, bottom), static (orange, middle) and degenerating (red, top) axons for WT and cdKO neurons at 1dpi. Classification is performed as follows: 1) regenerating axons extend past the site of injury, 2) degenerating axons either fully degenerate or show substantial beading and rupture and 3) static axons show minimal or no sign of beading but fail to extend past the site of injury. (G) Quantification of degenerating axons in the axonal compartment at 1dpi reveals a significant decrease in the percentage of degenerating axons in cdKO neurons. Data from 4 (E) or 6 (F-G) independent experiments, presented as mean ± SD (F-G) and bootstrap 95% CI versus WT(G). Student t-test (G). **p<0.01

**Figure S2:** Deletion of Pten and Socs3 enhances mitochondrial transport and morphology in regenerating axons. (A) Pairwise comparison of mitochondrial moving frequency between WT and cdKO axons at each timepoint and condition shows a comparable moving frequency between naive WT and cdKO axons, as well as at 0dpi and at 1dpi in static axons. On the other hand, regenerating cdKO axons at 1dpi reveal a higher mitochondrial moving frequency as compared to their WT counterparts. (B) Pairwise comparison of anterograde-only mitochondrial moving frequency between WT and cdKO axons at each timepoint and condition reveals a comparable moving frequency between naive WT and cdKO axons, as well as at 0dpi and at 1dpi in static axons. Conversely, regenerating cdKO axons at 1dpi reveal a higher anterograde mitochondrial moving frequency as compared to their WT counterparts. (C) Pairwise comparison of mitochondrial length between WT and cdKO axons at each timepoint and condition discloses a comparable size of mitochondria between naive WT and cdKO axons, as well as at 0dpi and at 1dpi in static axons. On the contrary, regenerating cdKO axons at 1dpi show a higher mitochondrial length as compared to WT ones. (D) Pairwise comparison of mitochondrial occupancy between WT and cdKO axons at each timepoint and condition shows no significant difference in axonal mitochondrial mass between WT and cdKO axons in any condition or timepoint. Data from 4 independent experiments, presented as mean ± SD (A-C) or median ± 25-75^th^CI (D) and bootstrap 95% CI versus relative naive. Welch (A-C) and Kruskal-Wallis ANOVA (D). ***p<0.001, ****p<0.0001

**Figure S3:** Deletion of Pten and Socs3 increases axonal energy production and glycolysis after injury. (A-B) Comparison of raw axonal ATP measurements identifies no significant difference in the intra-axonal concentration of ATP in naive axons between WT and cdKO neurons (A). Normalized measurements reveal a significantly higher intra-axonal concentration of ATP in cdKO axons as compared to WT right after injury at 0dpi and at 1dpi, in both static and regenerating axons (B). Datapoints were normalized against the median of their relative naive uninjured control per independent experiment. (C-D) Comparison of raw axonal lactate measurements identifies no significant difference in the intra-axonal concentration of lactate in naive axons between WT and cdKO neurons (C). Normalized measurements disclose a significantly higher intra-axonal concentration of lactate in cdKO axons as compared to WT right after axotomy at 0dpi and at 1dpi in regenerating axons (D). Datapoints were normalized against the median of their relative naive control per independent experiment. (E) Relative percentage representation of mitochondrial ATP (oligomycin-sensitive, green, top) and glycolytic ATP (oligomycin- and 2-deoxyglucose -sensitive, red, bottom) at 1dpi. Both static and regenerating cdKO axons show a higher relative percentage of glycolytic ATP as compared to WT axons. The percentage value was obtained by subtracting the baseline measurement in the presence of both 10 µM oligomycin and 50 mM 2-deoxyglucose from the measurements in normal medium and oligomycin-only. Then, the oligomycin-only measurement was expressed as a percentage of the measurement in normal medium. Data from 5 (A-B) or 4 (C-E) independent experiments, presented as median ± 25-75^th^ CI (A-D) and bootstrap 95% CI versus WT or mean ± SEM (E). Mann-Whitney U-test (A-D). **p<0.01, ***p<0.001, ****p<0.0001

**Figure S4:** Downregulation of axonal glycolysis reverses the regeneration phenotype of Pten and Socs3 deleted neurons. (A) Quantification of the percentage of degenerating axons at 1dpi shows that galactose treatment does not affect WT axons but reverses the reduced regeneration phenotype of cdKO axons. (B) Graphical representation of the percentages of degenerating (red, top), static (orange, middle) and regenerating (green, bottom) axons for WT and cdKO neurons. Galactose treatment reverses the phenotype of cdKO neurons. (C) Quantification of mitochondrial moving frequency at 1dpi shows a minor decrease in mitochondrial transport in static axons cultured in galactose as compared to glucose medium, but no significant decrease in regenerating axons. (D) Quantification of mitochondrial length in cdKO axons at 1dpi shows a minor but significant increase in mitochondrial elongation in regenerating axons when cultured in galactose. This is possibly due to compensatory mitochondrial fusion associated with increased oxidative phosphorylation. (E) Quantification of axonal mitochondrial occupancy in cdKO neurons at 1dpi shows no difference in mitochondrial density between axons cultured in glucose or galactose medium. Data from 4 independent experiments, presented as mean ± SD (A-B) or median ± 25-75^th^ CI (C-E) and bootstrap 95% CI versus glucose. Student t-test (A) or Mann-Whitney U-test (C-E). *p<0.05

## REFERENCES

1. Varadarajan, S.G., Hunyara, J.L., Hamilton, N.R., Kolodkin, A.L., and Huberman, A.D. (2022). Central nervous system regeneration. Cell 185, 77–94. 10.1016/j.cell.2021.10.029.

2. Fawcett, J.W. (2020). The Struggle to Make CNS Axons Regenerate: Why Has It Been so Difficult? Neurochem Res 45, 144–158. 10.1007/s11064-019-02844-y.

3. Sun, F., Park, K.K., Belin, S., Wang, D., Lu, T., Chen, G., Zhang, K., Yeung, C., Feng, G., Yankner, B.A., et al. (2011). Sustained axon regeneration induced by co-deletion of PTEN and SOCS3. Nature 480, 372–375. 10.1038/nature10594.

4. Jin, D., Liu, Y., Sun, F., Wang, X., Liu, X., and He, Z. (2015). Restoration of skilled locomotion by sprouting corticospinal axons induced by co-deletion of PTEN and SOCS3. Nature Communications 2015 6:1 *6*, 1–12. 10.1038/ncomms9074.

5. Cartoni, R., Norsworthy, M.W., Bei, F., Wang, C., Li, S., Zhang, Y., Gabel, C. V., Schwarz, T.L., and He, Z. (2016). The Mammalian-Specific Protein Armcx1 Regulates Mitochondrial Transport during Axon Regeneration. Neuron 92, 1294–1307. 10.1016/j.neuron.2016.10.060.

6. Cartoni, R., Pekkurnaz, G., Wang, C., Schwarz, T.L., and He, Z. (2017). A high mitochondrial transport rate characterizes CNS neurons with high axonal regeneration capacity. PLoS One 12, 1–12. 10.1371/journal.pone.0184672.

7. Zhou, B., Yu, P., Lin, M.Y., Sun, T., Chen, Y., and Sheng, Z.H. (2016). Facilitation of axon regeneration by enhancing mitochondrial transport and rescuing energy deficits. Journal of Cell Biology 214, 103–119. 10.1083/jcb.201605101.

8. Mar, F.M., Simões, A.R., Leite, S., Morgado, M.M., Santos, T.E., Rodrigo, I.S., Teixeira, C.A., Misgeld, T., and Sousa, M.M. (2014). CNS axons globally increase axonal transport after peripheral conditioning. Journal of Neuroscience 34, 5965–5970. 10.1523/JNEUROSCI.4680-13.2014.

9. Beckers, A., Masin, L., Van Dyck, A., Bergmans, S., Vanhunsel, S., Zhang, A., Verreet, T., Poulain, F.E., Farrow, K., and Moons, L. (2023). Optic nerve injury-induced regeneration in the adult zebrafish is accompanied by spatiotemporal changes in mitochondrial dynamics. Neural Regen Res 18, 219. 10.4103/1673-5374.344837.

10. Petrova, V., Nieuwenhuis, B., Fawcett, J.W., and Eva, R. (2021). Axonal organelles as molecular platforms for axon growth and regeneration after injury. Int J Mol Sci 22, 1–30. 10.3390/ijms22041798.

11. Hopkins, E.L., Gu, W., Kobe, B., and Coleman, M.P. (2021). A Novel NAD Signaling Mechanism in Axon Degeneration and its Relationship to Innate Immunity. Front Mol Biosci 8, 703532. 10.3389/FMOLB.2021.703532/BIBTEX.

12. Conforti, L., Gilley, J., and Coleman, M.P. (2014). Wallerian degeneration: an emerging axon death pathway linking injury and disease. Nature Reviews Neuroscience 2014 15:6 *15*, 394–409. 10.1038/nrn3680.

13. Lewis, T.L., Turi, G.F., Kwon, S.K., Losonczy, A., and Polleux, F. (2016). Progressive Decrease of Mitochondrial Motility during Maturation of Cortical Axons In Vitro and In Vivo. Current Biology 26, 2602–2608. 10.1016/j.cub.2016.07.064.

14. Taylor, A.M., Blurton-Jones, M., Rhee, S.W., Cribbs, D.H., Cotman, C.W., and Jeon, N.L. (2005). A microfluidic culture platform for CNS axonal injury, regeneration and transport. Nature Methods 2005 2:8 *2*, 599–605. 10.1038/nmeth777.

15. Rodriguez, A.R., de Sevilla Müller, L.P., and Brecha, N.C. (2014). The RNA binding protein RBPMS is a selective marker of ganglion cells in the mammalian retina. Journal of Comparative Neurology 522, 1411–1443. 10.1002/cne.23521.

16. Nadal-Nicolás, F.M., Galindo-Romero, C., Lucas-Ruiz, F., Marsh-Amstrong, N., Li, W., Vidal-Sanz, M., Agudo-Barriuso, M., Nadal-Nicolás, F.M., Galindo-Romero, C., Lucas-Ruiz, F., et al. (2023). Pan-retinal ganglion cell markers in mice, rats, and rhesus macaques. Zoological Research, 2023, Vol. 44, Issue 1, Pages: 226-248 *44*, 226–248. 10.24272/J.ISSN.2095-8137.2022.308.

17. Lewis, T.L., Kwon, S.-K., Lee, A., Shaw, R., and Polleux, F. (2018). MFF-dependent mitochondrial fission regulates presynaptic release and axon branching by limiting axonal mitochondria size. Nat Commun 9, 5008. 10.1038/s41467-018-07416-2.

18. Imamura, H., Huynh Nhat, K.P., Togawa, H., Saito, K., Iino, R., Kato-Yamada, Y., Nagai, T., and Noji, H. (2009). Visualization of ATP levels inside single living cells with fluorescence resonance energy transfer-based genetically encoded indicators. Proc Natl Acad Sci U S A 106, 15651–15656. 10.1073/pnas.0904764106.

19. Huang, N., Li, S., Xie, Y., Han, Q., Xu, X.M., and Sheng, Z.H. (2021). Reprogramming an energetic AKT-PAK5 axis boosts axon energy supply and facilitates neuron survival and regeneration after injury and ischemia. Current Biology 31, 3098–3114. 10.1016/j.cub.2021.04.079.

20. Heiden, M.G.V., Cantley, L.C., and Thompson, C.B. (2009). Understanding the warburg effect: The metabolic requirements of cell proliferation. Science (1979) 324, 1029–1033. 10.1126/SCIENCE.1160809/ASSET/C00CA2CE-14F9-4352-AC43-7D1BB04B41D8/ASSETS/GRAPHIC/324_1029_F4.JPEG.

21. Liberti, M. V., and Locasale, J.W. (2016). The Warburg Effect: How Does it Benefit Cancer Cells? Trends Biochem Sci 41, 211–218. 10.1016/j.tibs.2015.12.001.

22. Frey, P.A. (1996). The Leloir pathway: a mechanistic imperative for three enzymes to change the stereochemical configuration of a single carbon in galactose. The FASEB Journal 10, 461–470. 10.1096/FASEBJ.10.4.8647345.

23. Li, H., Guglielmetti, C., Sei, Y.J., Zilberter, M., Le Page, L.M., Shields, L., Yang, J., Nguyen, K., Tiret, B., Gao, X., et al. (2023). Neurons require glucose uptake and glycolysis in vivo. Cell Rep 42, 112335. 10.1016/J.CELREP.2023.112335.

24. Robinson, B.H., Petrova-Benedict, R., Buncic, J.R., and Wallace, D.C. (1992). Nonviability of cells with oxidative defects in galactose medium: A screening test for affected patient fibroblasts. Biochem Med Metab Biol 48, 122–126. 10.1016/0885-4505(92)90056-5.

25. Aguer, C., Gambarotta, D., Mailloux, R.J., Moffat, C., Dent, R., McPherson, R., and Harper, M.E. (2011). Galactose Enhances Oxidative Metabolism and Reveals Mitochondrial Dysfunction in Human Primary Muscle Cells. PLoS One 6, e28536. 10.1371/JOURNAL.PONE.0028536.

26. Cheng, X.-T., Huang, N., and Sheng, Z.-H. (2022). Programming axonal mitochondrial maintenance and bioenergetics in neurodegeneration and regeneration. 10.1016/j.neuron.2022.03.015.

27. Hopkins, E.L., Gu, W., Kobe, B., and Coleman, M.P. (2021). A Novel NAD Signaling Mechanism in Axon Degeneration and its Relationship to Innate Immunity. Front Mol Biosci 8, 703532. 10.3389/FMOLB.2021.703532/BIBTEX.

28. Conforti, L., Gilley, J., and Coleman, M.P. (2014). Wallerian degeneration: an emerging axon death pathway linking injury and disease. Nature Reviews Neuroscience 2014 15:6 *15*, 394–409. 10.1038/nrn3680.

29. Zheng, X., Boyer, L., Jin, M., Mertens, J., Kim, Y., Ma, L., Ma, L., Hamm, M., Gage, F.H., and Hunter, T. (2016). Metabolic reprogramming during neuronal differentiation from aerobic glycolysis to neuronal oxidative phosphorylation. Elife 5. 10.7554/eLife.13374.

30. Iwata, R., Casimir, P., Erkol, E., Boubakar, L., Planque, M., López, I.M.G., Ditkowska, M., Gaspariunaite, V., Beckers, S., Remans, D., et al. (2023). Mitochondria metabolism sets the species-specific tempo of neuronal development. Science (1979). 10.1126/SCIENCE.ABN4705.

31. Bonvento, G., and Bolaños, J.P. (2021). Astrocyte-neuron metabolic cooperation shapes brain activity. Cell Metab 33, 1546–1564. 10.1016/J.CMET.2021.07.006.

32. Chamberlain, K.A., and Sheng, Z.H. (2019). Mechanisms for the maintenance and regulation of axonal energy supply. J Neurosci Res 97, 897–913. 10.1002/JNR.24411.

33. Wei, Y., Miao, Q., Zhang, Q., Mao, S., Li, M., Xu, X., Xia, X., Wei, K., Fan, Y., Zheng, X., et al. (2023). Aerobic glycolysis is the predominant means of glucose metabolism in neuronal somata, which protects against oxidative damage. Nat Neurosci 26, 2081– 2089. 10.1038/s41593-023-01476-4.

34. Wolfe, A.D., Koberstein, J.N., Smith, C.B., Stewart, M.L., Gonzalez, I.J., Hammarlund, M., Hyman, A.A., Stork, P.J.S., Goodman, R.H., and Colón-Ramos, D.A. (2024). Local and dynamic regulation of neuronal glycolysis in vivo. Proceedings of the National Academy of Sciences 121, e2314699121. 10.1073/pnas.2314699121.

35. Díaz-García, C.M., Mongeon, R., Lahmann, C., Koveal, D., Zucker, H., and Yellen, G. (2017). Neuronal Stimulation Triggers Neuronal Glycolysis and Not Lactate Uptake. Cell Metab 26, 361–374.e4. 10.1016/j.cmet.2017.06.021.

36. Zala, D., Hinckelmann, M.V., Yu, H., Lyra Da Cunha, M.M., Liot, G., Cordelières, F.P., Marco, S., and Saudou, F. (2013). Vesicular glycolysis provides on-board energy for fast axonal transport. Cell 152, 479–491. 10.1016/j.cell.2012.12.029.

37. Hinckelmann, M.V., Virlogeux, A., Niehage, C., Poujol, C., Choquet, D., Hoflack, B., Zala, D., and Saudou, F. (2016). Self-propelling vesicles define glycolysis as the minimal energy machinery for neuronal transport. Nat Commun 7, 1–13. 10.1038/ncomms13233.

38. Mc Cluskey, M., Dubouchaud, H., Nicot, A., and Saudou, F. (2023). A vesicular Warburg effect: Aerobic glycolysis occurs on axonal vesicles for local NAD+ recycling and transport. Traffic. 10.1111/tra.12926.

39. Ketschek, A., Sainath, R., Holland, S., and Gallo, G. (2021). The axonal glycolytic pathway contributes to sensory axon extension and growth cone dynamics. Journal of Neuroscience 41, 6637–6651. 10.1523/JNEUROSCI.0321-21.2021.

40. Santos, R., Lokmane, L., Ozdemir, D., Traoré, C., Agesilas, A., Hakibilen, C., Lenkei, Z., and Zala, D. (2023). Local glycolysis fuels actomyosin contraction during axonal retraction. Journal of Cell Biology 222. 10.1083/JCB.202206133.

41. Hubley, M.J., Locke, B.R., and Moerland, T.S. (1996). The effects of temperature, pH, and magnesium on the diffusion coefficient of ATP in solutions of physiological ionic strength. Biochimica et Biophysica Acta (BBA) - General Subjects 1291, 115–121. 10.1016/0304-4165(96)00053-0.

42. Takenaka, T., Ohnishi, Y., Yamamoto, M., Setoyama, D., and Kishima, H. (2023). Glycolytic System in Axons Supplement Decreased ATP Levels after Axotomy of the Peripheral Nerve. eNeuro 10. 10.1523/ENEURO.0353-22.2023.

43. Li, F., Sami, A., Noristani, H.N., Slattery, K., Qiu, J., Groves, T., Wang, S., Veerasammy, K., Chen, Y.X., Morales, J., et al. (2020). Glial Metabolic Rewiring Promotes Axon Regeneration and Functional Recovery in the Central Nervous System. Cell Metab 32, 767–785.e7. 10.1016/j.cmet.2020.08.015.

44. Spinelli, J.B., and Haigis, M.C. (2018). The multifaceted contributions of mitochondria to cellular metabolism. Nat Cell Biol 20, 745–754. 10.1038/s41556-018-0124-1.

45. Rangaraju, V., Lewis, T.L., Hirabayashi, Y., Bergami, M., Motori, E., Cartoni, R., Kwon, S.K., and Courchet, J. (2019). Pleiotropic Mitochondria: The Influence of Mitochondria on Neuronal Development and Disease. Journal of Neuroscience 39, 8200–8208. 10.1523/JNEUROSCI.1157-19.2019.

46. Tracey, T.J., Steyn, F.J., Wolvetang, E.J., and Ngo, S.T. (2018). Neuronal lipid metabolism: Multiple pathways driving functional outcomes in health and disease. Front Mol Neurosci 11, 331939. 10.3389/FNMOL.2018.00010/BIBTEX.

47. Xing, J., Lukomska, A., Rheaume, B.A., Kim, J., Sajid, M.S., Damania, A., and Trakhtenberg, E.F. (2023). Post-injury born oligodendrocytes incorporate into the glial scar and contribute to the inhibition of axon regeneration. Development 150. 10.1242/DEV.201311.

48. Crabtree, H.G. (1935). The differential effect of radium radiation on the carbohydrate metabolism of normal and tumour tissues irradiated at low temperature. Biochem J 29, 2334–2343. 10.1042/BJ0292334.

49. Leibinger, M., Müller, A., Gobrecht, P., Diekmann, H., Andreadaki, A., and Fischer, D. (2013). Interleukin-6 contributes to CNS axon regeneration upon inflammatory stimulation. Cell Death Dis 4, e609. 10.1038/cddis.2013.126.

50. Petrova, V., Pearson, C.S., Ching, J., Tribble, J.R., Solano, A.G., Yang, Y., Love, F.M., Watt, R.J., Osborne, A., Reid, E., et al. (2020). Protrudin functions from the endoplasmic reticulum to support axon regeneration in the adult CNS. Nat Commun 11, 1–15. 10.1038/s41467-020-19436-y.

51. Schindelin, J., Arganda-Carreras, I., Frise, E., Kaynig, V., Longair, M., Pietzsch, T., Preibisch, S., Rueden, C., Saalfeld, S., Schmid, B., et al. (2012). Fiji: an open-source platform for biological-image analysis. Nature Methods 2012 9:7 *9*, 676–682. 10.1038/nmeth.2019.

52. Ho, J., Tumkaya, T., Aryal, S., Choi, H., and Claridge-Chang, A. (2019). Moving beyond P values: data analysis with estimation graphics. Nature Methods 2019 16:7 *16*, 565–566. 10.1038/s41592-019-0470-3.

53. Masin, L., Claes, M., Bergmans, S., Cools, L., Andries, L., Davis, B.M., Moons, L., and De Groef, L. (2021). A novel retinal ganglion cell quantification tool based on deep learning. Scientific Reports 2021 11:1 *11*, 1–13. 10.1038/s41598-020-80308-y.

54. Yasukawa, H., Ohishi, M., Mori, H., Murakami, M., Chinen, T., Aki, D., Hanada, T., Takeda, K., Akira, S., Hoshijima, M., et al. (2003). IL-6 induces an anti-inflammatory response in the absence of SOCS3 in macrophages. Nat Immunol 4, 551–556. 10.1038/ni938.

55. Lesche, R., Groszer, M., Gao, J., Wang, Y., Messing, A., Sun, H., Liu, X., and Wu, H. (2002). Cre/loxP-mediated inactivation of the murine Pten tumor suppressor gene. Genesis 32, 148–149. 10.1002/GENE.10036.

56. Pereiro, X., Ruzafa, N., Acera, A., Fonollosa, A., David Rodriguez, F., and Vecino, E. (2018). Dexamethasone protects retinal ganglion cells but not Müller glia against hyperglycemia in vitro. PLoS One 13, 1–17. 10.1371/journal.pone.0207913.

57. Basu, H., Ding, L., Pekkurnaz, G., Cronin, M., and Schwarz, T.L. (2020). Kymolyzer, a Semi-Autonomous Kymography Tool to Analyze Intracellular Motility. Curr Protoc Cell Biol 87. 10.1002/CPCB.107.

58. Ho, J., Tumkaya, T., Aryal, S., Choi, H., and Claridge-Chang, A. (2019). Moving beyond P values: data analysis with estimation graphics. Nature Methods 2019 16:7 *16*, 565–566. 10.1038/s41592-019-0470-3.

