## Supplemental figures for "Local glycolysis supports injury-induced axonal regeneration"

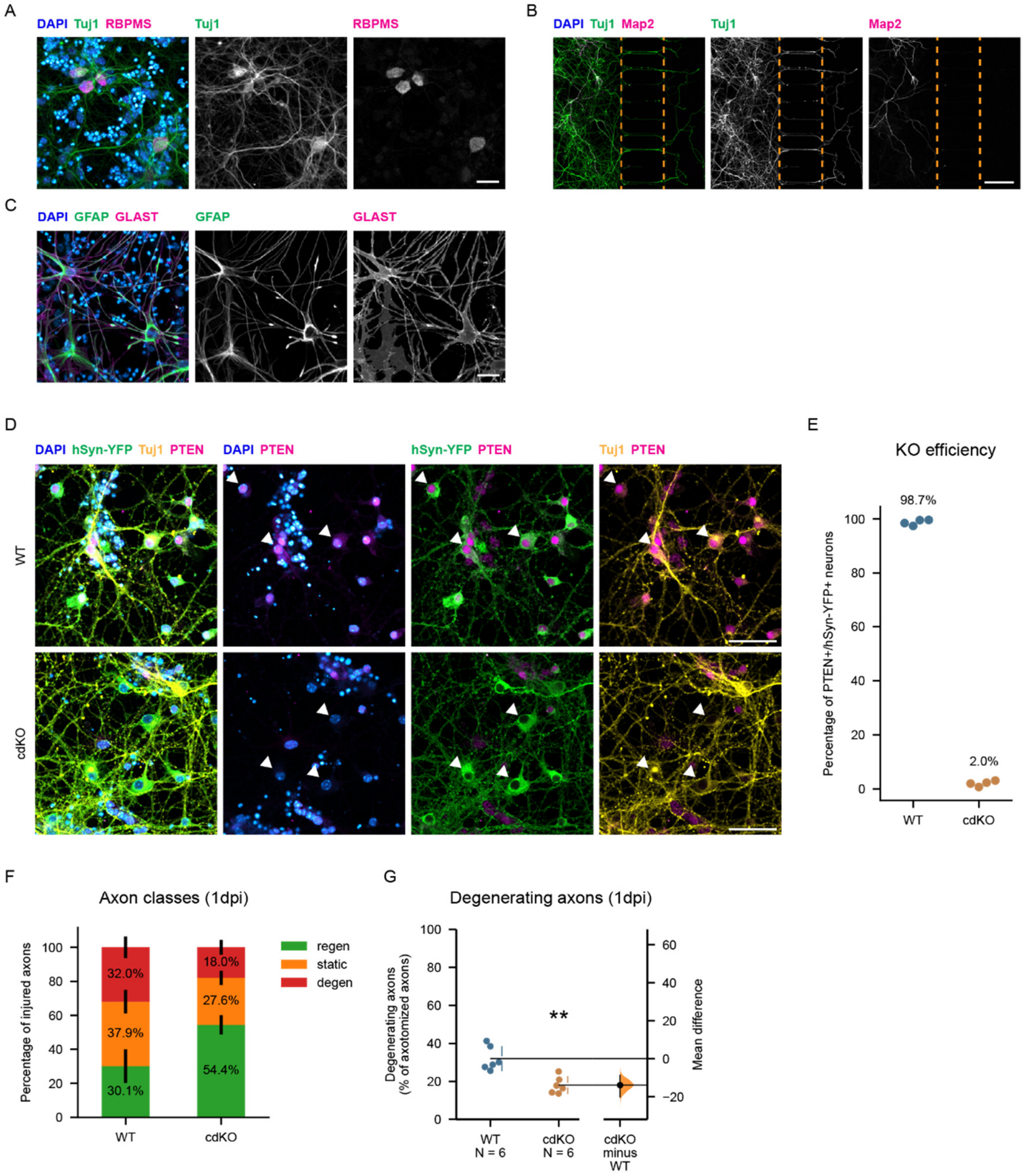


***Figure S1: Validation of the retinal cell culture and microfluidic model.***

(A) Immunocytochemistry for Tuj1 and RBPMS at 14 days *in vitro* (DIV) identifies retinal ganglion cells (RGCs), co-labelled by both markers, in the mixed retinal cell culture. Scale bar 25 μm.

(B) Immunocytochemistry for Tuj1 and Map2 at 14DIV identifies Tuj1+ RGCs in the somatic compartment and their axons in the axonal one. Importantly, the dendritic marker Map2 is restricted to the somatic compartment. The edges of the microgrooves are delineated by the orange dotted line. Scale bar 100 μm.

(C) Immunocytochemistry for GFAP and GLAST at 14DIV revels the presence of retinal macroglia, co-labelled by both markers. Scale bar 25 μm

(D) Immunolabelling for PTEN and Tuj1 shows specific loss of PTEN within Tuj1+/hSyn-YFP+ RGCs upon transduction with an AAV2/2-Syn-Cre viral vector (cdKO) as compared to

WT cells, indicating specific recombination and gene knock-out. hSyn-YFP is encoded by *AAV2/2-hSyn1-ATeam$^{YEMK}$-WPRE-hGHp*, see Methods. Scale bar 50 µm.

(E) Quantification of PTEN signal in the somata of hSyn-YFP+ RGCs showes that on average 98.7% of the RGCs are positive for PTEN at 14DIV in the WT condition as compared to only 2% in the cdKO culture, indicating that an efficient knock-out is achieved. (hSyn-YFP encoded by *AAV2/2-hSyn1-ATeam$^{YEMK}$-WPRE-hGHp*)

(F) Graphical representation of the percentages of regenerating (green, bottom), static (orange, middle) and degenerating (red, top) axons for WT and cdKO neurons at 1dpi. Classification is performed as follows: 1) regenerating axons extend past the site of injury, 2) degenerating axons either fully degenerate or show substantial beading and rupture and 3) static axons show minimal or no sign of beading but fail to extend past the site of injury.

(G) Quantification of degenerating axons in the axonal compartment at 1dpi reveals a significant decrease in the percentage of degenerating axons in cdKO neurons.

Data from 4 (E) or 6 (F-G) independent experiments, presented as mean ± SD (F-G) and bootstrap 95% CI versus WT(G). Student t-test (G). **$p<0.01$

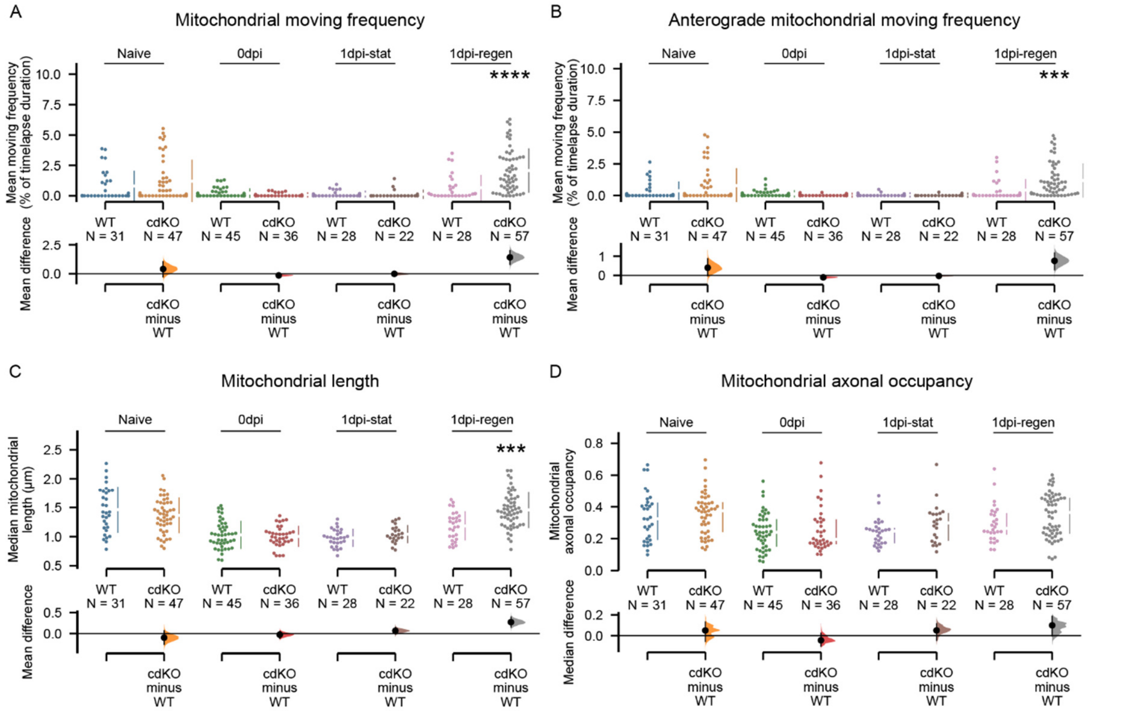


***Figure S2: Deletion of Pten and Socs3 enhances mitochondrial transport and morphology in regenerating axons.***

(A) Pairwise comparison of mitochondrial moving frequency between WT and cdKO axons at each timepoint and condition shows a comparable moving frequency between naive WT and cdKO axons, as well as at 0dpi and at 1dpi in static axons. On the other hand, regenerating cdKO axons at 1dpi reveal a higher mitochondrial moving frequency as compared to their WT counterparts.

(B) Pairwise comparison of anterograde-only mitochondrial moving frequency between WT and cdKO axons at each timepoint and condition reveals a comparable moving frequency between naive WT and cdKO axons, as well as at 0dpi and at 1dpi in static axons. Conversely, regenerating cdKO axons at 1dpi reveal a higher anterograde mitochondrial moving frequency as compared to their WT counterparts.

(C) Pairwise comparison of mitochondrial length between WT and cdKO axons at each timepoint and condition discloses a comparable size of mitochondria between naive WT and cdKO axons, as well as at 0dpi and at 1dpi in static axons. On the contrary, regenerating cdKO axons at 1dpi show a higher mitochondrial length as compared to WT ones.

(D) Pairwise comparison of mitochondrial occupancy between WT and cdKO axons at each timepoint and condition shows no significant difference in axonal mitochondrial mass between WT and cdKO axons in any condition or timepoint.

Data from 4 independent experiments, presented as mean ± SD (A-C) or median ± 25-75$^{th}$CI (D) and bootstrap 95% CI versus relative naive. Welch (A-C) and Kruskal-Wallis ANOVA (D). ***p<0.001, ****p<0.0001

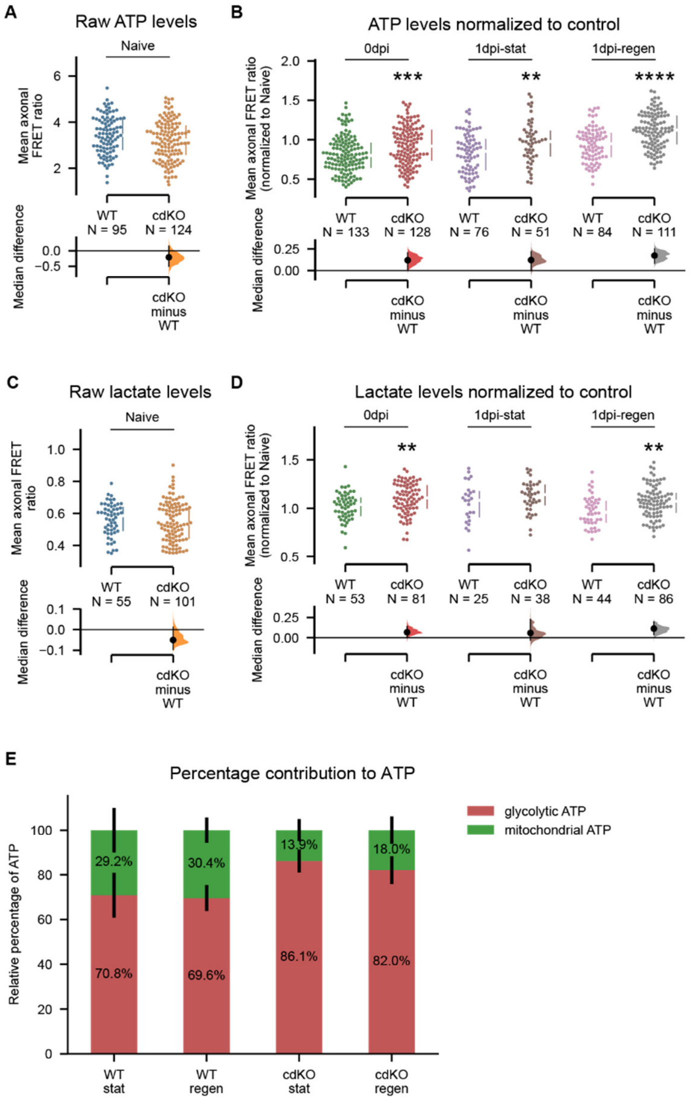


***Figure S3: Deletion of Pten and Socs3 increases axonal energy production and glycolysis after injury.***

(A-B) Comparison of raw axonal ATP measurements identifies no significant difference in the intra-axonal concentration of ATP in naive axons between WT and cdKO neurons (A). Normalized measurements reveal a significantly higher intra-axonal concentration of ATP in cdKO axons as compared to WT right after injury at 0dpi and at 1dpi, in both static and regenerating axons (B). Datapoints were normalized against the median of their relative naive uninjured control per independent experiment.

(C-D) Comparison of raw axonal lactate measurements identifies no significant difference in the intra-axonal concentration of lactate in naive axons between WT and cdKO neurons (C). Normalized measurements disclose a significantly higher intra-axonal concentration of lactate in cdKO axons as compared to WT right after axotomy at 0dpi and at 1dpi in regenerating axons (D). Datapoints were normalized against the median of their relative naive control per independent experiment.

(E) Relative percentage representation of mitochondrial ATP (oligomycin-sensitive, green, top) and glycolytic ATP (oligomycin- and 2-deoxyglucose -sensitive, red, bottom) at 1dpi.

Both static and regenerating cdKO axons show a higher relative percentage of glycolytic ATP as compared to WT axons. The percentage value was obtained by subtracting the baseline measurement in the presence of both 10 µM oligomycin and 50 mM 2-deoxyglucose from the measurements in normal medium and oligomycin-only. Then, the oligomycin-only measurement was expressed as a percentage of the measurement in normal medium.

Data from 5 (A-B) or 4 (C-E) independent experiments, presented as median ± 25-75$^{th}$ CI (A-D) and bootstrap 95% CI versus WT or mean ± SEM (E). Mann-Whitney U-test (A-D). **p<0.01, ***p<0.001, ****p<0.0001

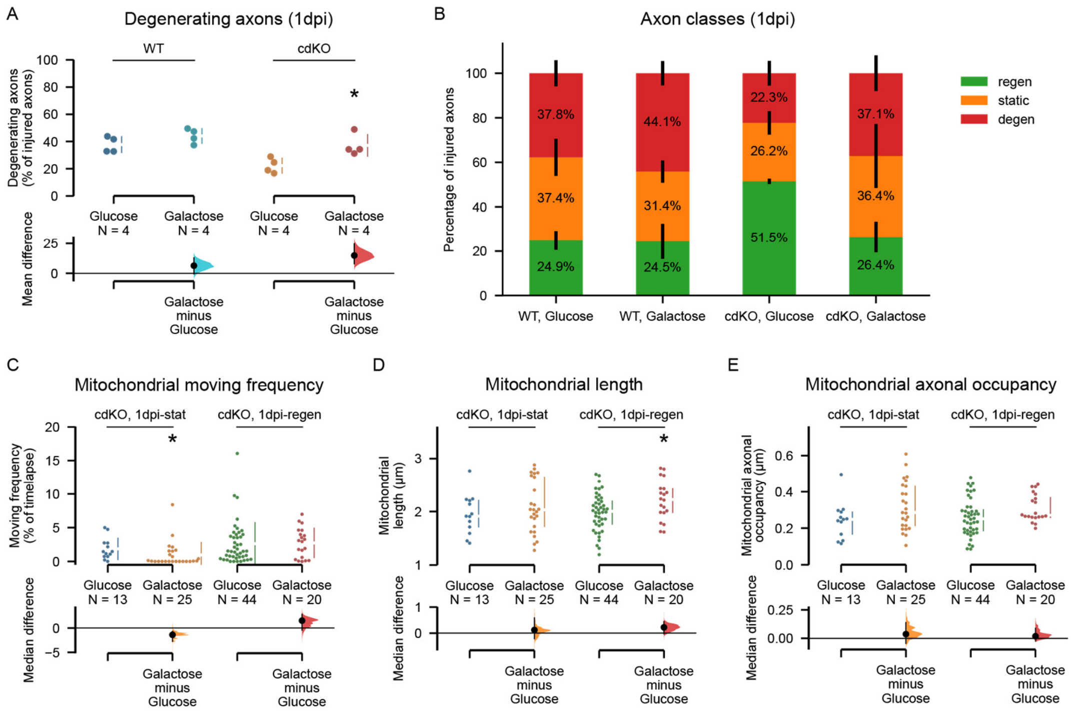


***Figure S4: Downregulation of axonal glycolysis reverses the regeneration phenotype of Pten and Socs3 deleted neurons.***

(A) Quantification of the percentage of degenerating axons at 1dpi shows that galactose treatment does not affect WT axons but reverses the reduced regeneration phenotype of cdKO axons.

(B) Graphical representation of the percentages of degenerating (red, top), static (orange, middle) and regenerating (green, bottom) axons for WT and cdKO neurons. Galactose treatment reverses the phenotype of cdKO neurons.

(C) Quantification of mitochondrial moving frequency at 1dpi shows a minor decrease in mitochondrial transport in static axons cultured in galactose as compared to glucose medium, but no significant decrease in regenerating axons.

(D) Quantification of mitochondrial length in cdKO axons at 1dpi shows a minor but significant increase in mitochondrial elongation in regenerating axons when cultured in galactose. This is possibly due to compensatory mitochondrial fusion associated with increased oxidative phosphorylation.

(E) Quantification of axonal mitochondrial occupancy in cdKO neurons at 1dpi shows no difference in mitochondrial density between axons cultured in glucose or galactose medium.

Data from 4 independent experiments, presented as mean ± SD (A-B) or median ± 25-75$^{th}$ CI (C-E) and bootstrap 95% CI versus glucose. Student t-test (A) or Mann-Whitney U-test (C-E). *$p<0.05$
